# Targeted *DUX4* base editing improves muscle function in an iPSC-derived model of childhood-onset FSHD

**DOI:** 10.64898/2026.07.28.741079

**Authors:** Peter J. Houweling, Vanessa G. Crossman, Leonit Kiriaev, Kathrin Mattes, Chrystal Tiong, Chantal A. Coles, Claire Hogan, Natasha Tuano, Richard J Mills, Sara E. Howden, Katy de Valle, Kathryn N. North, Ian R Woodcock

## Abstract

Facioscapulohumeral muscular dystrophy (FSHD) is one of the most common dominant muscular dystrophies and remains without an approved disease modifying therapy. Caused by the aberrant expression of the cytotoxic gene *DUX4,* FSHD is typically diagnosed in adulthood, however clinical onset in children (<18 years of age) is often associated with a more severe and rapid disease. While clinical trials are underway, a lack of human-specific pre-clinical models limit effective testing of potential therapies, particularly in children. To fill this gap, we describe here the development of induced pluripotent stem cell-derived 2-and 3-dimensional skeletal muscle models of children with clinically defined mild, moderate, and severe FSHD. These iPSC-derived muscle models replicate key features of FSHD, including aberrant *DUX4* mRNA expression, muscle atrophy, and weakness, which correlate with the individuals’ specific disease severity. Next, we assessed the efficacy of adenine base editing (ABE) as a potential gene therapy approach to treat FSHD. *DUX4*-targeted ABE reduced *DUX4* mRNA expression, improved muscle area and force generation in the most severe individual. Together this work supports the use of iPSC-derived skeletal muscle models as a less invasive method to study childhood-onset FSHD and establishes targeted *DUX4* gene editing therapies as a potential treatment approach.

## Introduction

Facioscapulohumeral muscular dystrophy (FSHD) is an autosomal dominant neuromuscular disorder caused by the aberrant expression of the normally repressed transcription factor double homeobox 4 (*DUX4*), which leads to progressive skeletal muscle atrophy and weakness. With an estimated prevalence of ∼1 per 8 000 individuals worldwide, FSHD is the third most common muscular dystrophy [1]. Although typically diagnosed in adults [2], approximately 30% of FSHD cases present in childhood (<18 years of age) and correspond with a more rapid and severe disease progression [3]. Childhood-onset FSHD often represents the severe end of the FSHD disease pathology spectrum which poses a disproportionate clinical burden and a unique opportunity to study disease pathology in individuals with the most rapidly evolving disease trajectory.

At a molecular level, this clinical heterogeneity is underpinned by a well-characterised genetic mechanism. The underlying genetic cause of FSHD is driven by the epigenetic dysregulation of the D4Z4 macrosatellite repeat array on chromosome 4q35A. This can occur via a contracted D4Z4 repeat array to <10 units, as in FSHD type 1, or via specific mutations in chromatin modifier genes (e.g *SMCHD1, DNMT3B* or *LRIF1*), as in FSHD type 2 [4, 5]. Both scenarios result in the hypomethylation of chromosome 4 and the aberrant expression of *DUX4* in somatic cells [6]. In skeletal muscle the re-expression of *DUX4* is cytotoxic and results in atrophy and weakness [7].

The discovery that *DUX4* is the genetic driver of FSHD enabled the development of various cell and animal models. However, the *DUX4* gene is primate specific, with mice not containing the equivalent D4Z4 repeat array or an endogenous *DUX4* transcript [8, 9]. Although the mouse ortholog, *Dux,* has similar functions with its over-expression resulting in pathology [10], most pre-clinical mouse models rely on the artificial overexpression of human *DUX4*, which only partially mimics the molecular and pathological features of FSHD [11]. In parallel, primary muscle biopsies and isolated skeletal muscle cell cultures obtained from individuals with FSHD show sporadic *DUX4* expression and the activation of downstream cytotoxic gene expression signals [12, 13]. However, limitations in cell proliferation capacity and challenges in obtaining muscle biopsies, particularly from children, has restricted the use of these human-specific models to study FSHD. Furthermore, heterogeneity in disease presentation and progression confounds investigations into FSHD specific disease mechanisms, negatively impacting clinical trial design and cohort stratification, and hampering our ability to translate potential targeted therapies for FSHD.

These limitations highlight the need for human-specific, scalable models that more accurately capture the genetic and epigenetic context of FSHD. To address these challenges, induced pluripotent stem cells (iPSCs) have emerged as a promising tool to generate human-specific tissues that overcome many of the limitations of traditional model systems. In FSHD, iPSC-derived skeletal muscle progenitors and terminally differentiated 2-dimensional (D) multinucleated myotube cultures have been shown to recapitulate key disease-relevant features. This includes retention of the D4Z4 repeat contraction and permissive epigenetic context [14], as well as transcriptional dysregulation, including aberrant *DUX4* expression and activation of downstream target genes (such as *ZSCAN4* and *TRIM43*), muscle apoptosis and cell death [15].

While 2D culture systems capture key molecular features, more advanced 3D engineered muscle models provide a higher level of physiological relevance. 3D engineered skeletal muscle models generated from the iPSCs of adults with mosaic FSHD also recapitulate core disease features, including *DUX4* mRNA expression, reduced myofiber diameter, impaired sarcomere organization, and compromised contractile function [16]. Notably, Franken *et al.* demonstrated that several therapeutic candidates which show promising improvements in 2D muscle cultures, show limited benefit, and in some cases are detrimental, when tested in more complex 3D muscle models [16].

These findings underscore both the potential and limitations of current pre-clinical FSHD models and highlight the need for clinically defined physiologically relevant human skeletal muscle models that better represent disease pathology. To date, no preclinical (primary or iPSC-derived skeletal muscle models) of childhood-onset FSHD have been described, nor have such models been used to evaluate potential therapeutic candidates *in vitro*, a crucial step before potential treatments can be tested.

In this study, we describe the first iPSC-derived skeletal muscle models of individuals with childhood-onset FSHD that span the spectrum of disease severity (mild, moderate and severe). Using both 2-and 3D engineered muscle tissues we evaluate FSHD associated phenotypes *in vitro* including *DUX4* mRNA expression, muscle volume and strength. Furthermore, in the most severely affected individuals’ iPSCs we apply adenine base editing (ABE) to target *DUX4* gene expression and examine the efficacy of this gene-editing approach to improve FSHD pathology. Collectively, these findings establish iPSC-derived skeletal muscle as a clinically relevant platform for modelling disease severity in childhood-onset FSHD and provide strong support for the continued development of DUX4-targeted gene editing therapies.

## Results

### Individuals with childhood-onset FSHD show the spectrum of disease severity

Four individuals (1 female (F): 3 male (M)) with a genetically confirmed diagnosis of childhood-onset FSHD type 1 were selected from the Australian pediatric FSHD longitudinal outcomes study (iFSHD-LOS, (ACTRN12621001293853)). D4Z4 repeat size was confirmed during diagnosis, with two individuals containing 5 repeats (FSHD_1 and 6) and two containing 3 repeats (FSHD_7 and 8) (Table 1). Blood samples were collected at the time of clinical assessment, with iPSC lines generated from the peripheral blood mononuclear cells (PBMCs) of each individual (Table 1).

**Table 1:** FSHD patient information including study number, sex (Female (F) / Male (M)), age of symptom onset and sample collection, genetic confirmation, repeat/D4Z4 size, FSHD clinical score summary (mild, moderate and severe).

| Study number | Sex | Age of symptom onset (years) | Age at sample collection (years) | Genetic confirmation | Repeat/D4Z4 size | FSHD Clinical score (0 – 15) |
| --- | --- | --- | --- | --- | --- | --- |
| FSHD_1 | F | <10 | 16 | FSHD1 | 5 | 4 - mild/mod |
| FSHD_6 | M | <10 | 19 | FSHD1 | 5 | 2 – mild |
| FSHD_7 | M | <10 | 13 | FSHD1 | 3 and 6 | 7 – mod |
| FSHD_8 | M | <10 | 17 | FSHD1 | 3 | 15 – severe |

Clinical severity was determined using three functional assessments: the FSHD Clinical Score (FCS), pediatric FSHD-Composite Outcome Measure (FSHD-COM Peds), and 6-minute walk distance (6MWD).

Participants selected for iPSC generation ensured the full phenotypic spectrum was represented with two participants (FSHD_1 and FSHD_6) having mild disease as represented by an FCS severity score of 4/15 and 2/15 respectively, one participant with moderate disease (FSHD_7) with FCS 7/15 and another with severe disease (FSHD_8) with FCS 14/15. (Fig. 2A).

The FSHD-COM Peds total scores demonstrate similar disease severity, with FSHD_1 (18/84) and FSHD_6 (4/84) showing mild symptoms, FSHD_7 (33/84) moderate and FSHD_8 the most severe (77/84) functional decline (Fig. 2B).

6MWD measures walking capacity, with reduced distance correlating with greater disease severity. Mildly affected participants FSHD_1 and FSHD_6 walked ∼650–700Rm, and the moderately affected participant FSHD_7 walked ∼400Rm. The most severely affected participant, FSHD_8, was non ambulant (0Rm) at the time of sample collection (Fig.R2C).

The detailed functional assessment performed for each individual replicate the association between D4Z4 repeat size and disease severity, with FSHD_1 and 6 (5 D4Z4 repeats) showing a mild phenotype and FSHD_7 and FSHD_8 (3 D4Z4 repeats) a moderate and severe pathology (Table 1). We next sought to determine if the same clinical phenotype can be replicated *in vitro* using iPSC-derived 2-and 3D skeletal muscle models.

### iPSC lines generated from four individuals with childhood-onset FSHD are pluripotent but do not express *DUX4* mRNA

iPSC lines generated from the PBMCs of our selected FSHD individual’s (FSHD_1 (MCHTB396), 6 (MCHTB405), 7 (MCHTB406) and 8 (MCHTB411)) were next confirmed to be pluripotent. iPSCs were generated using Sendai virus carrying the reprogramming factors POU5F1 (OCT4), SOX2, KLF4 and MYC as previously published ([17, 18], Supp Fig. 1A). Each line expressed pluripotency markers OCT4, TRA-1-60, SSEA-4 and TRA-1-81 by flow cytometry (Supp Fig. 1B). Differentiation to the three germ cell layers (ectoderm, mesoderm and endoderm) was performed using confirmed pluripotency (Supp Fig. 1C). The generation of an ectoderm cell lineage was confirmed by the expression of Nestin and PAX6 (Supp Fig. 1D). Mesoderm was confirmed by the downregulation of *OCT4* and up-regulation of *Brachyury* and *CXCR4* mRNA and differentiation to endoderm was confirmed by the expression of SOX17 (Supp Fig. 1E).

No differences in the iPSC replication rate were observed between each iPSC lines or selected healthy control (Supp Fig. 1G). Furthermore, *DUX4* mRNA was not detected in any of the FSHD or healthy control iPSCs (Supp Fig. H). After confirming pluripotency, we differentiated these iPSCs into skeletal muscle progenitors and generated both 2 and 3D engineered muscle tissue models.

**Graphical Abstract/Figure 1:**
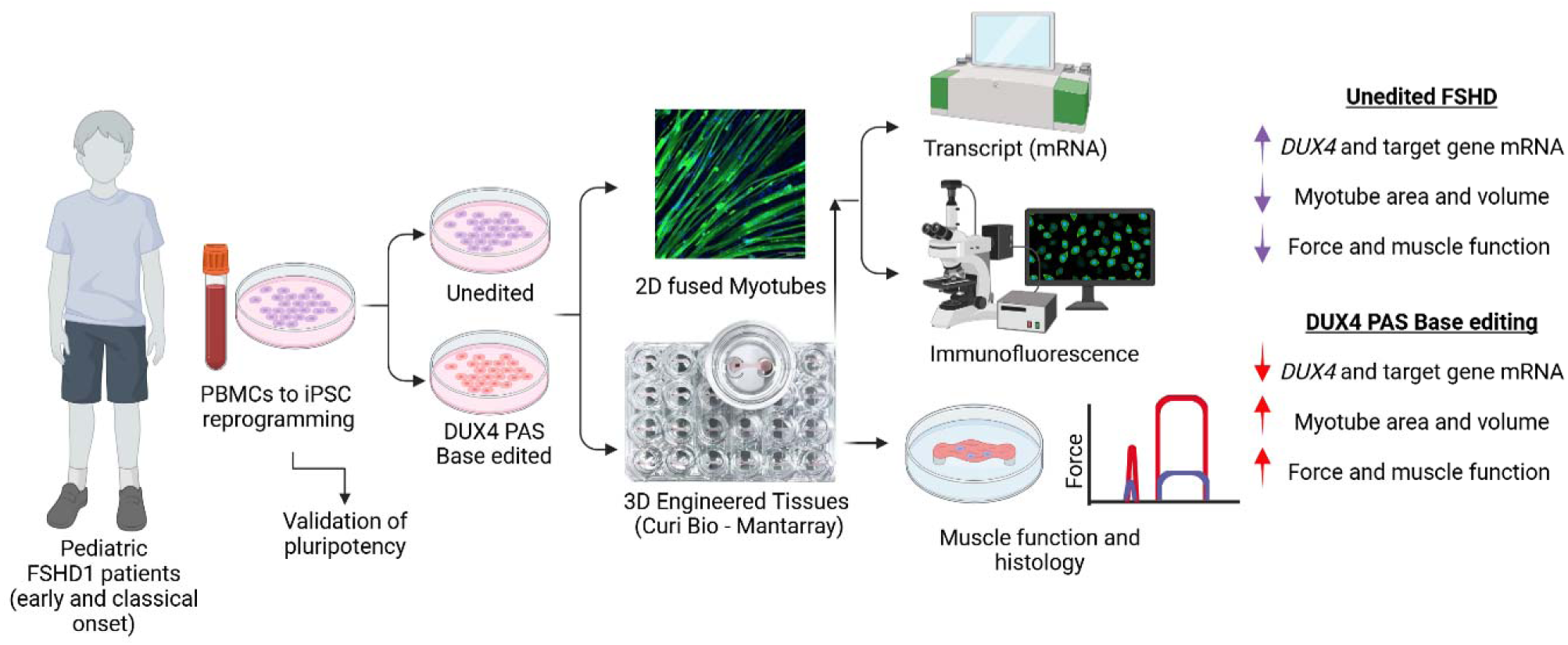
Induced pluripotent stem cell (iPSC) derived 2-and 3-dimensional skeletal muscle models generated from four individuals with genetically confirmed childhood-onset facioscapulohumeral muscular dystrophy type 1 demonstrate key pathological features of disease including aberrant DUX4 mRNA and downstream target gene expression, reduced muscle area, volume and force, all without the need of an invasive muscle biopsy. Adenine base editing (ABE) of the DUX4 polyadenylation sequence (PAS) reduced DUX4 mRNA expression and improved muscle area, volume and force, providing important proof-of-concept data for the use of targeted gene editing approaches for FSHD.

**Figure 2:**
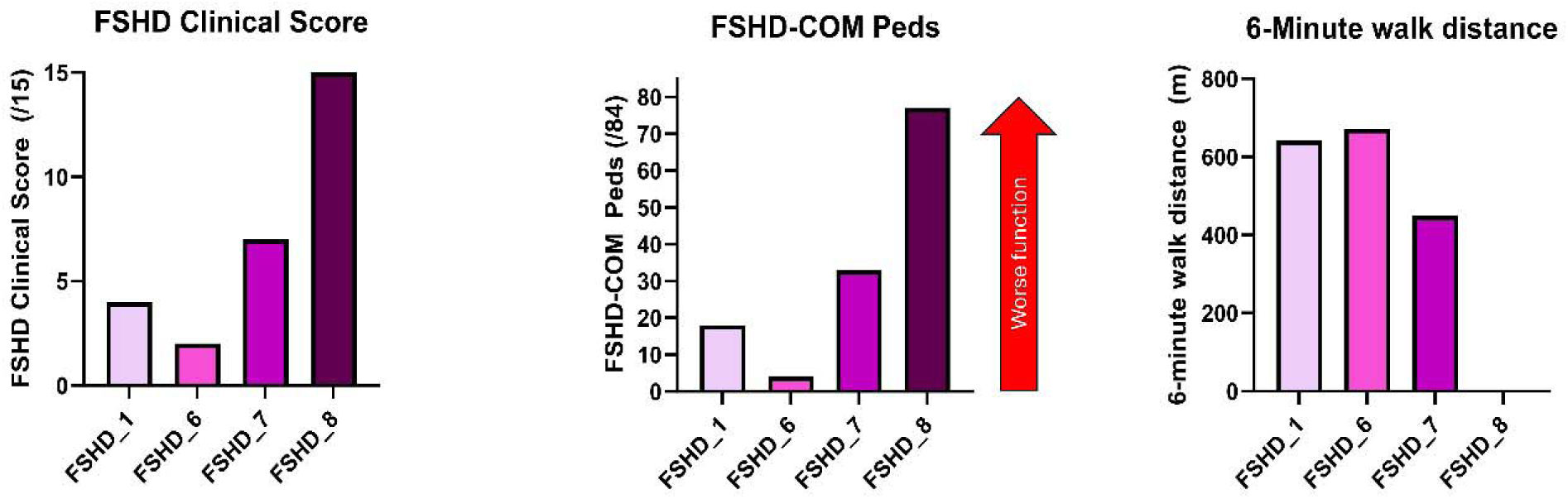
Childhood-onset FSHD clinical phenotyping: A) FSHD Clinical score, B) FSHD-COM Peds and C) 6-Minute walk distance data show patient specific disease severity outcomes. At the time of blood collection and iPSC generation, FSHD individuals 1 and 6 (with 5 D4Z4 repeats) show more mild symptoms compared to FSHD individuals 7 and 8 (with 3 D4Z4 repeats) that show a more moderate (FSHD_7) and severe (FSHD_8) phenotype.

### 2D iPSC-derived skeletal muscle myotubes show temporal *DUX4* expression and morphological changes that correlate with disease severity

We first performed a time course analysis of myogenic differentiation using iPSC derived 2D skeletal muscle cells, progressing from myoblasts (day 0) to multinucleated myotubes at days 3, 4, 5, and 6. Across this differentiation series, key molecular and morphological features of FSHD were quantified (Fig. 3A).

**Figure 3:**
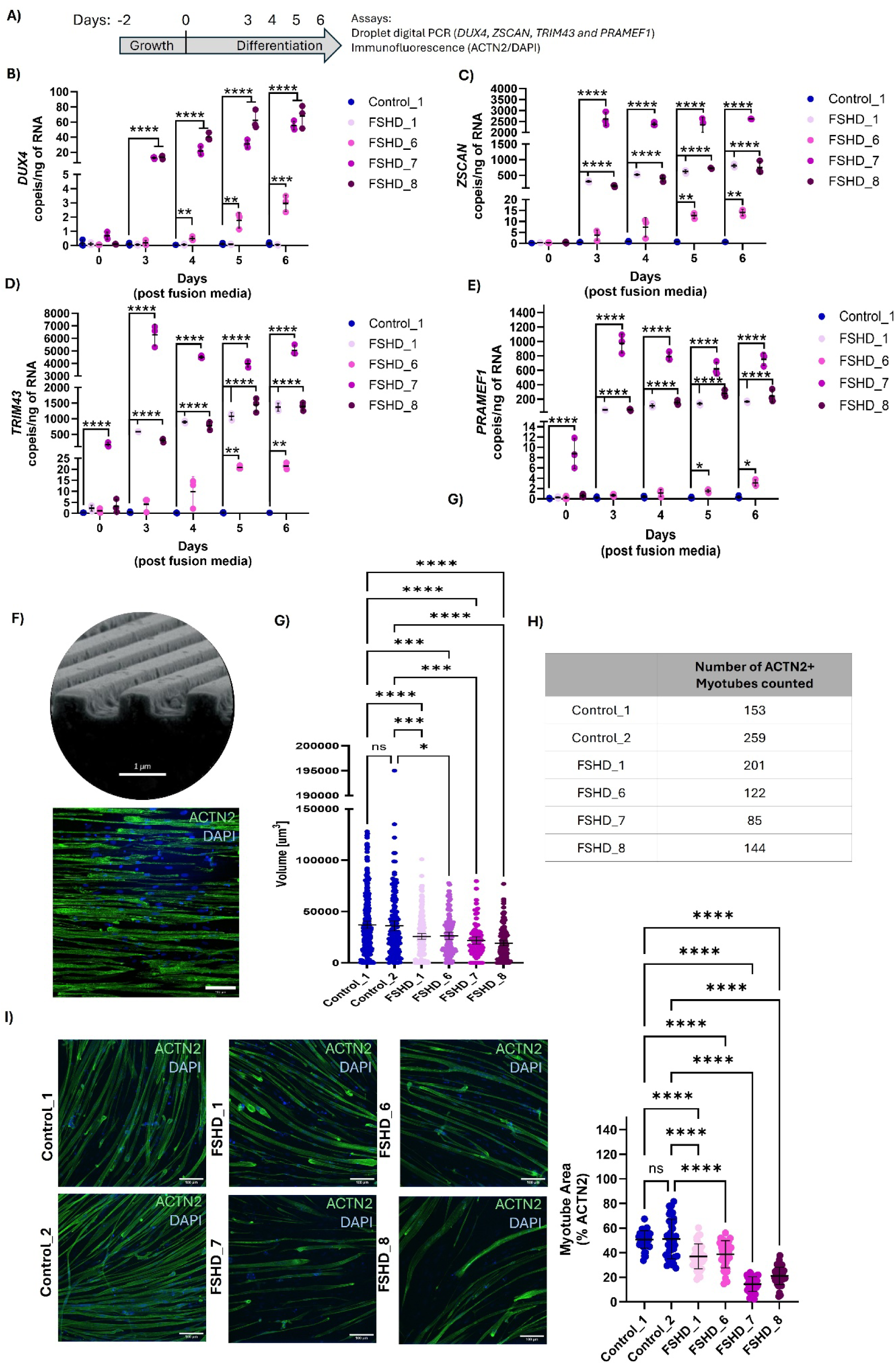
2D iPSC-derived FSHD muscles express DUX4 and downstream target genes, are thinner and show reduced myotube area which correlates with clinical severity; A) Experimental timeline: myoblasts were cultured for two days in growth media, followed by differentiation in fusion media for up to six days. Droplet digital PCR (ddPCR) was used to quantify expression of B) DUX4, C) ZSCAN4, D) TRIM43, and E) PRAMEF1. F) Immunofluorescence assessed ACTN2-positive myotubes. G) Volume of ACTN2+ myotubes were determined based on H) individual myotubes counted. ACTN2 positive area (I) were assessed in healthy controls (n = 2) and FSHD (n = 4). ddPCR analyses were determined based on N = 3 replicates / individual / timepoint (B – E). Myotube volume was determined based on the number of myotubes shown in (G) and ACTN2+ Area (I) was determined based on three independent analyses of between 36 and 45 images / well. The mean of individual datapoints is shown + standard deviation (SD). All statistical analyses were performed using a one-way ANOVA with corrections for multiple testing (Tukey). P-value *<0.05; ** <0.01, *** <0.001, **** <0.0001.

Droplet digital PCR (ddPCR) confirmed significant increases in *DUX4* mRNA expression in three (FSHD_6, 7 and 8) of four FSHD lines, with upregulation detectable as early as day 3 in the two more severe FSHD individuals (Fig. 3B). No *DUX4* mRNA was detected in the healthy control (Fig. 3B).

Consistent with FSHD pathology in primary skeletal muscles, increased expression of the downstream *DUX4*-trageted genes *ZSCAN4*, *TRIM43* and *PRAMEF1* occurred in all iPSC-derived myotubes, with the highest levels detected in the more severe individuals (Fig. 3C, D and E).

To evaluate myofiber morphology, myotubes were grown on nano etched tissue culture plates and imaged using the sarcomeric protein α actinin 2 (ACTN2) as a marker of myotube volume and area (Fig. 3F). ACTN2 positive (+) myotubes were assessed (Fig. 3G) with all FSHD lines exhibiting a reduction in myotube volume relative to controls, with the greatest decline observed in FSHD_7 and FSHD_8 (Fig. 3H). Similarly, compared to healthy controls ACTN2+ myotube area was reduced across all FSHD individuals (Control 1 = 51% ± 7.2 SD, Control 2 = 51% ± 16 SD, FSHD_1 = 37% ± 10 SD, FSHD_6 39% ± 11 SD, FSHD_7 = 14% ± 6 SD, FSHD_8 = 21% ± 7 SD) with the most pronounced deficits corresponding to those with more severe disease (Fig. 3I).

### 3D engineered muscle tissues of childhood-onset FSHD replicate disease pathology with reduced muscle force corresponding to disease severity

Next, we evaluated FSHD-associated phenotypes in more complex 3D engineered muscle tissue (EMT) environment, assessing targeted transcript activation, contractile function, and myofiber architecture after 14 days in culture (Fig. 4A).

**Figure 4:**
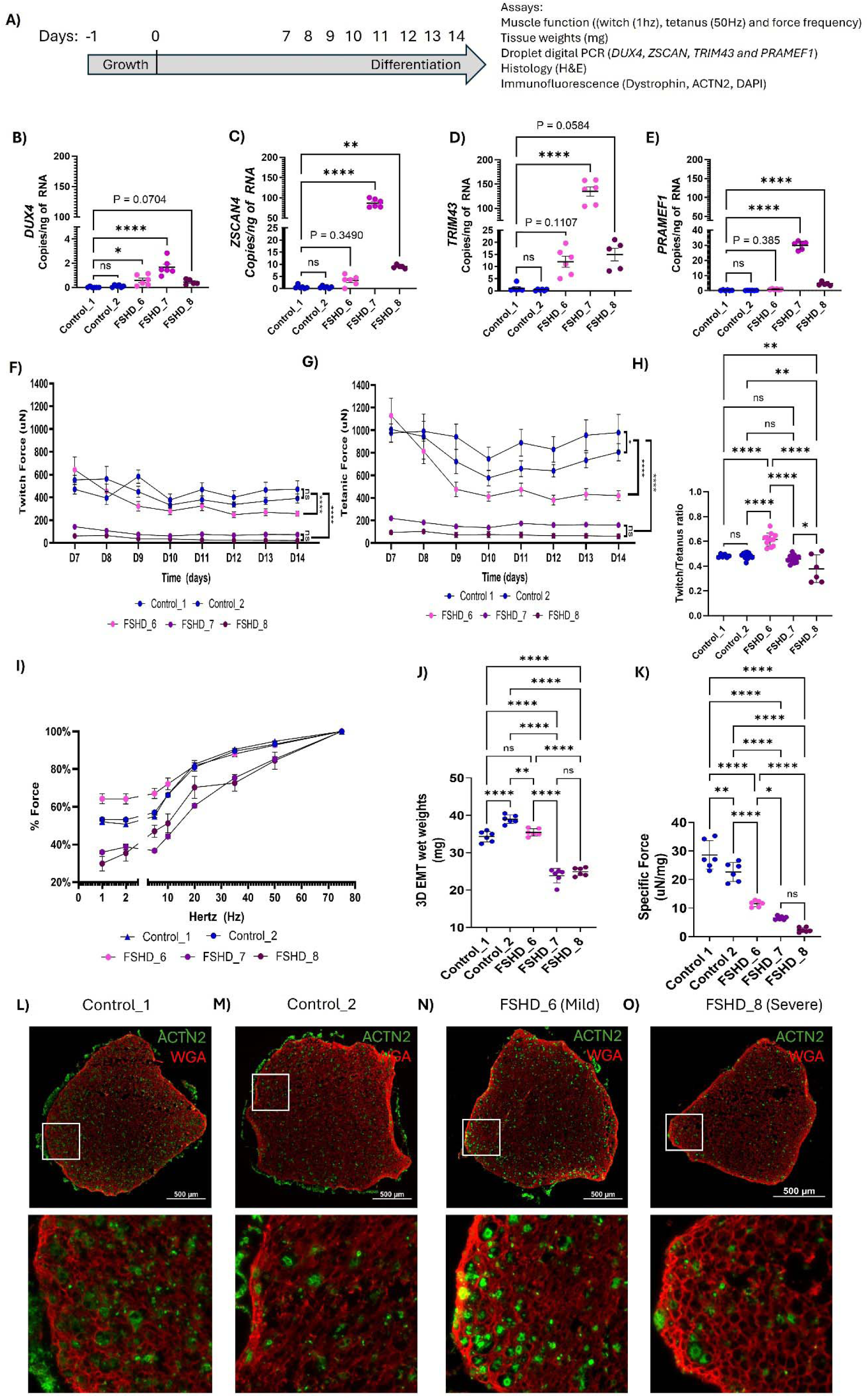
3D EMTs show increased DUX4 and downstream target gene expression with reduced force correlating with patient-specific disease severity. A) Experimental timeline: myoblasts were cultured for 24hrs in growth media, followed by differentiation in fusion media for up to 14 days. B) DUX4 mRNA levels show significant increases in FSHD_6 and 7, and a trend (P = 0.07) in FSHD_8. Downstream genes C) ZSCAN4, D) TRIM43, and E) PRAMEF1 were increased in FSHD_7 and 8, compared to healthy controls. Longitudinal functional assessments from day 7 (D7) in culture were measured daily for twitch (F) and tetanus (G) until day 14 (D14) with significant reductions in force seen in all FSHD lines, compared to healthy controls (Two-way ANOVA). H) Twitch:Tetanus ratios were significantly different between FSHD and control 3D EMTs at 14 days. I) Normalized force frequency analyses were also different between FSHD and control. Individual 3D EMT tissue weights show reduced mass for FSHD_7 and 8, and specific force (µN/mg) were significantly reduced in FSHD_6, 7 and 8 compared to controls. L-M) Immunohistochemistry imaging of ACTN2 (green) and wheat-germ (red) show the presence of myotubes in all cultures with less ACTN2+ myotubes seen in more severe EMTs, compared to control and mild lines. The mean of individual datapoints is shown, + standard deviation (SD). All statistical analyses were performed using ANOVA (one or two way) with corrections for multiple testing (Tukey). N = 6 3D EMTs per individual were analysed for all measures. P-value *<0.05; ** <0.01, *** <0.001, **** <0.0001.

Similar to 2D, mRNA analysis confirmed the expression of *DUX4* and downstream targets *ZSCAN4*, *TRIM43* and *PRAMEF1* in 3D EMT from individuals with FSHD. *DUX4* mRNA was significantly elevated in 2 out of 3 FSHD individuals, compared to healthy controls (FSHD_6 and 7, Fig. 4B), with increases in *ZSCAN4* (Fig. 4C), *TRIM43* (Fig.4D), and *PRAMEF1* (Fig.4E) mRNA, observed in the more severe individuals (FSHD_7 and 8).

Contractile assessments from day 7 in culture revealed marked deficits in muscle strength with both twitch (1 Hz) and tetanic (50Hz) forces reduced in FSHD 3D EMTs, compared to controls (Fig. 4F and G). Compared to healthy controls the mildly affected individual (FSHD_6) exhibited a significant increase in twitch (Control_1 = P<0.001; Control_2 = P <0.001) and tetanic force (Control_1 = P<0.05; Control_2 = P=0.0592) at day 7; however, force progressively declined until stabilizing at ∼50% reduction by day 9 to 14 (P<0.0001). In contrast, the more severely affected individuals (FSHD_7 and FSHD_8) showed substantial force deficits from day 7 onward, with a ∼95% reduction in both twitch (Fig. 4F, P<0.0001) and tetanic (Fig. 4G, P<0.0001) forces, relative to healthy controls from day 7 to 14 (Two-way ANOVA data summary Supplementary Table 1 and 2).

As a measure of maturation, twitch:tetanus ratios showed no differences between the two control lines, with the highest ratio seen in mild (FSHD_6) compared to both severe FSHD lines (FSHD_7 and 8, Fig. 4H) which suggests a more immature muscle phenotype in those with more severe FSHD. At day 14, force-frequency curves revealed that healthy controls generate substantially higher absolute forces across all stimulation frequencies, whereas severe FSHD EMTs display a blunted contractile response (Fig. 4I). Consistent with this functional decline, 3D EMT wet weight were reduced in all individuals with FSHD, with the most severely affected showing the greatest decline (Fig. 4J). When normalized to tissue mass, specific force remained low in all FSHD 3D EMTs, with the greatest deficit observed in the more severe individuals (FSHD_7 and FSHD_8, Fig. 4K). Finally, immunohistochemical analyses of 3D FSHD EMTs suggested reduced ACTN2+ myofiber abundance in the severe individual compared to healthy controls and mild individuals (Fig. 4L – O).

### Adenine base editing suppresses *DUX4* and improves myotube morphology in 2D cultures

Adenine base editing (ABE) of *DUX4* has been proposed as a potential approach to attenuate the cytotoxic FSHD transcriptional program [19, 20]. However, the utility of ABE in 2 and 3D EMT models had not yet been evaluated. Therefore, we used a targeted ABE editing strategy and the iPSCs from the most severe individual (FSHD_8) to test the utility of this therapeutic strategy *in vitro* (Fig. 5A).

**Figure 5:**
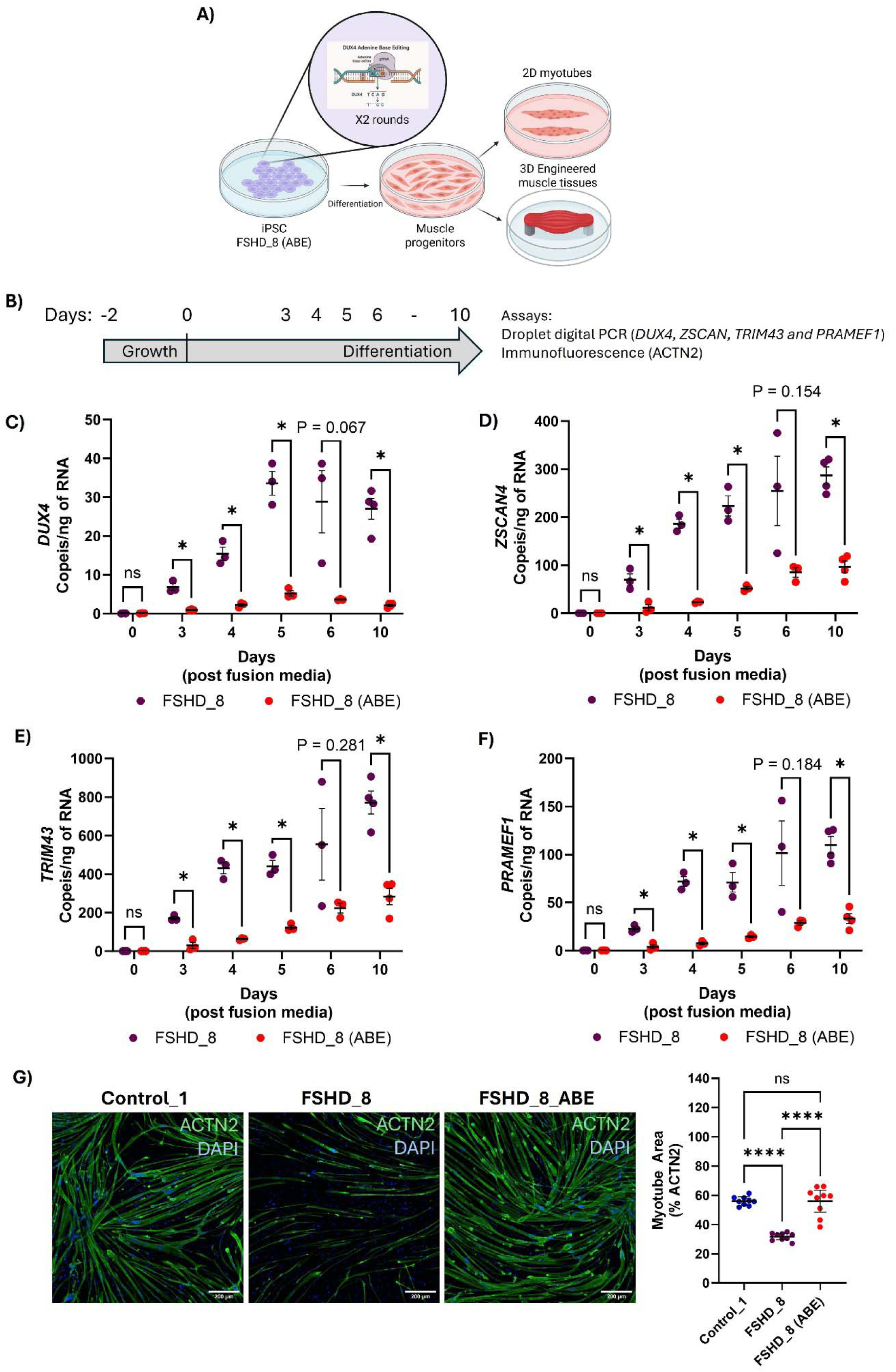
DUX4 adenine base editing reduces DUX4 mRNA and downstream target gene expression in 2D. A) Adine base editing strategy in iPSCs from FSHD_8. B) Experimental timeline: myoblasts that reached 100% confluency were transferred to fusion media for up to 10 days. C) DUX4 mRNA expression is significant reduction from at least day 3 post fusion media following ABE. D) ZSCAN4, E) TRIM43, and F) PRAMEF1 mRNA levels were all reduced compared to unedited myotubes (n = 3 replicates per individual). G) Improved muscle architecture was also observed following ABE, compared to unedited (FSHD_8) myotubes with the % ACTN2 positive myotube area no longer different between ABE and healthy control muscles (n = 9 images per individual). The mean of individual datapoints is shown, + standard deviation (SD). All statistical analyses were performed using one way ANOVA with corrections for multiple testing (Tukey). P-value *<0.05; ** <0.01, *** <0.001, **** <0.0001.

A time-course analyses of 2D myotubes from day 3 to 10 showed that the unedited FSHD line (FSHD_8) exhibited elevated *DUX4* mRNA during differentiation, whereas the ABE edited myotubes (FSHD_8 (ABE)) showed a reduction in *DUX4* mRNA from day 3 (Fig. 5C). In parallel, the expression of *DUX4* target genes *ZSCAN4*, *TRIM43*, and *PRAMEF1* was reduced in the ABE iPSC-derived muscles relative to unedited control (Fig. 5D–F).

We next assessed ACTN2+ myotube area to determine whether *DUX4* mRNA suppression translates to a structural improvement in FSHD myotubes. Compared with healthy controls, unedited FSHD_8 myotubes displayed reduced ACTN2+ myotube area (Control = 56% ± 3 SD, FSHD_8 = 32% ± 3 SD). In contrast, ABE resulted in a significant increase in ACTN2+ myotube area (FSHD_8_ABE = 56% ± 10 SD), restoring it to the same level observed in healthy controls (Fig. 5H).

### ABE partially rescues *DUX4*-associated molecular signatures and contractile function in 3D EMTs

We next tested whether ABE-mediated suppression of *DUX4* translated into functional improvements in 3D EMTs that were matured over an extended 24-day period (Fig. 6A).

**Figure 6:**
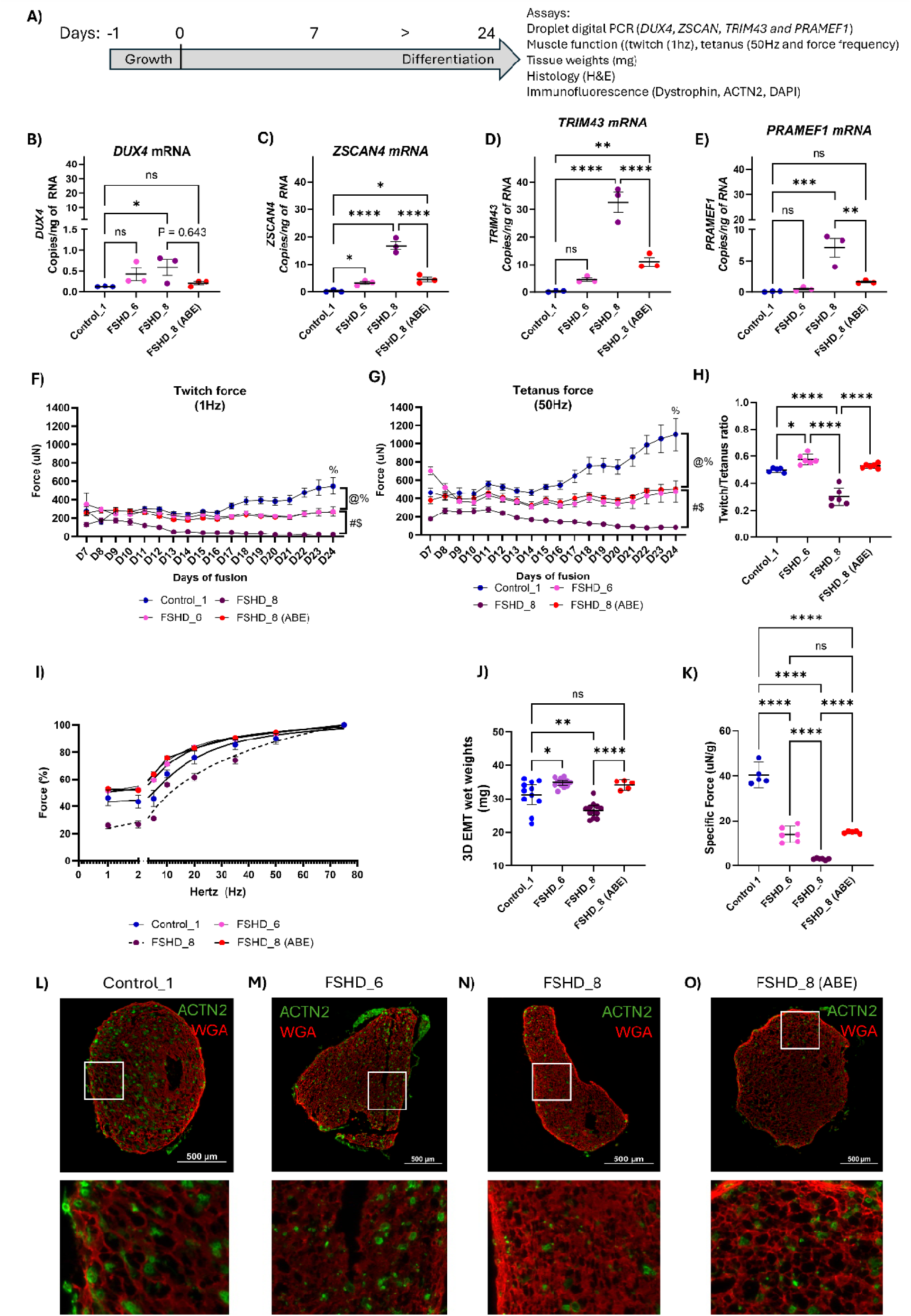
3D Muscle function improves following DUX4 base-editing A) Experimental timeline with assays performed. Longitudinal force data was collected daily from day 7 to 24, with endpoint analyses performed on EMT’s collected at day 24. B) DUX4 mRNA expression is reduced in ABE muscles, so that the expression was no longer different to controls at day 24, C) ZSCAN4, D) TRIM43 and E) PRAMEF1 mRNA levels were all reduced in ABE tissues, compared to unedited EMT’s, however the expression levels remained elevated compared to healthy controls for ZSCAN4 and TRIM43 (n = 3 EMTs per individual). F) Twitch force and G) Tetanic force were significantly reduced in all FSHD EMTs with ABE significantly improving muscle force compared to unedited EMTs at day 24 (Two-way ANOVA, n = 6 3D EMTs per individual, % = significant differences between control and FSHD_8, @ = significant difference between control and FSHD_6 and % = FSHD_8 (ABE), # = significant differences between FSHD_8 and FSHD6, $ = significant differences between FSHD_8 and FSHD_8 (ABE). H) Twitch:Tetanus ratio increased following ABE, compared to unedited EMTs with ABE 3D EMTs no longer different to controls. The mean of individual datapoints is shown, + standard deviation (SD). All statistical analyses were performed using ANOVA (one or two way) with corrections for multiple testing (Tukey). P-value *<0.05; ** <0.01, *** <0.001, **** <0.0001.

In matured 3D EMTs, unedited muscles (FSHD_8) exhibited elevated expression of *DUX4* (Fig. 6B) and downstream target genes *ZSCAN4*, *TRIM43* and *PRAMEF1*, compared with healthy controls (Fig. 6C–E). Compared to unedited controls, ABE resulted in a non-significant reduction in *DUX4* mRNA (P = 0.643), however *DUX4* mRNA levels were no longer significantly elevated compared to healthy controls (P>0.05, Fig. 6B). Similarly, the expression of downstream gene targets *ZSCAN4*, *TRIM43*, *PRAMEF1* were all significantly reduced with ABE, however both *ZSCAN4* (P<0.05) and *TRIM43* (P<0.01) mRNA levels remained elevated when compared to the healthy control (Fig. 6C–E).

Longitudinal contractility measurements demonstrated improved muscle function in ABE *DUX4* 3D EMTs. From day 7 to 10 ABE 3D EMTs generated similar twitch (1 Hz, Fig. 6F) and tetanic (50 Hz, Fig. 6H) force when compared to healthy controls (P>0.05). However, this improved function was not maintained as twitch and tetanic force increased in healthy controls over the 24-day analysis period (P<0.0001).

At 24 days both twitch and tetanic force outputs were reduced in untreated mild (FSHD_6, Twitch = 272 µN + 51 µN SD; Tetanus = 477 µN + 108 µN SD, P<0.0001) and severe (FSHD_8, Twitch = 26 µN + 2 µN SD; Tetanus = 88 µN + 5 µN SD, P<0.0001) EMTs compared with healthy controls (Twitch = 548 µN + 72 µN SD; Tetanus = 1099 µN + 141 µN SD, P<0.0001).

Compared to unedited EMTs, ABE generated significantly higher twitch (D24 FSHD_8 (ABE) = 268 µN ± 10 µN SD, P<0.0001) and tetanic (D24 FSHD_8 (ABE) = 506 µN ± 4 µN SD, P<0.0001) forces. From day 9 to 24 ABE 3D EMTs were indistinguishable from the mild FSHD constructs (D9 to D24 P>0.05, Supplementary Table 3 and 4).

Consistent with improved contractile output, twitch:tetanus ratio revealed improved maturation, with ABE shifting 3D EMTs towards healthy control and mild, relative to the unedited lines (Fig. 6H). Similarly, force–frequency analysis showed that mild and ABE EMTs produced greater absolute forces across stimulation frequencies, while unedited severe 3D EMTs exhibited a blunted response. ABE displayed an improved force–frequency profile relative to the unedited line, which were more consistent with the mild individual (Fig. 6I).

3D EMTs wet tissue weights were reduced in severe FSHD 3D EMTs (26.55mg ± 2.23 SD) compared with controls (31.16 mg ± 4.45 SD, P<0.001) and mild FSHD (34.8 mg ± 1.34 SD, Fig. 6J, P<0.05). ABE increased tissue mass (34.16 mg ± 1.33 SD, Fig. 6J) relative to unedited FSHD_8 (P<0.0001). Importantly, normalization of tetanic force to tissue mass showed that specific force remained significantly impaired in unedited mild and severe FSHD tissues, whereas ABE increased specific force, reaching the same level as the mild FSHD individual, but remaining significantly reduced compared to the healthy control (Fig. 6K, P<0.0001). Structural analysis of tissue cross-sections also showed that, compared to controls, ABE 3D EMTs showed improved structure relative to unedited, including increased abundance of ACTN2+ myofibers, which was consistent with improvements in muscle function unedited FSHD EMTs displayed reduced myofiber density (Fig. 6L–N).

## Discussion

This study explored the molecular, structural and functional consequences of childhood-onset FSHD using complementary 2 and 3D iPSC-derived skeletal muscle models. Using these models we demonstrate the potential efficacy of *DUX4* gene-targeted adenine base editing to mitigate disease-relevant phenotypes and improve muscle function in FSHD.

Using defined differentiation timelines in both 2D myotubes and 3D EMTs, we show that *DUX4* and its canonical downstream target genes (*ZSCAN4*, *TRIM43*, and *PRAMEF1*) are aberrantly activated in childhood-onset FSHD during skeletal muscle differentiation, with the level of *DUX4* (and downstream target gene) activation highest in more severely affected FSHD iPSC-derived muscle culture models. This data corresponds to a shorter D4Z4 repeat length and more advanced disease-specific pathobiology seen in these individuals. These findings reinforce a dose-dependent relationship between D4Z4 repeat size, *DUX4* activity, and disease severity.

The increase in *DUX4* mRNA coincides with defects in myotube formation and maturation, associated with reduced myotube volume and fewer ACTN2+ myotubes in 2D. This also culminates in markedly reduced twitch and tetanus forces in 3D EMTs. Again, the deficit in muscle force corresponded with the individual’s disease severity at the time of iPSC line generation, with 3D EMTs generated from the mild FSHD individual showing a 50% drop in muscle function over both 14 and 24 days in culture, compared to an almost 95% reduction in force in the more severe FSHD individuals. Together, these data establish iPSC-derived skeletal muscle models of childhood-onset FSHD as a viable tool to dissect the impact of disease severity and the pathogenic role of *DUX4* activation, impaired myogenesis, and compromised contractile outputs, without the need of an invasive muscle biopsy.

In severe FSHD, *DUX4* mRNA increased in 2D muscle cultures at the onset of muscle fusion (from Day 3 in culture) and remained detectable throughout the differentiation window, while DUX4-target genes mirrored this activation, but at a higher level of expression. *DUX4* expression is often described as “sporadic” and “low” in human primary muscle cultures and biopsies; our results replicate this and show low/no *DUX4* mRNA expression in individuals with mild disease, and greater levels in individuals with more severe disease. However, intermittent *DUX4* activation seen in mild patients is sufficient to drive pathology *in vitro*, with cumulative impacts on structural and functional measures seen in both 2 and 3D muscle cultures from individuals with both mild and severe FSHD. The concordant upregulation of *ZSCAN4*, *TRIM43*, and *PRAMEF1* mRNA in all FSHD muscle models supports the concept of a *DUX4*-centered transcriptional cascade that promotes disease pathology and drives severity.

Structural analyses of 2D myotube area and volume as well as the more complex 3D EMTs reflect these molecular findings with thinner, and less abundant myotubes across all FSHD lines, relative to healthy controls. Reduced numbers of ACTN2+ myotubes, and lower muscle mass in 3D EMTs support impaired muscle function and suggest reduced survival of skeletal muscle *in vitro,* which correlates with disease severity.

Furthermore, functional measurements of 3D EMTs provide a longitudinal analysis of muscle strength that correlates with an individual’s specific disease severity. Reduced twitch and tetanic force observed in both mild and severe FSHD 3D EMTs, with a greater decline in specific force in severe FSHD supports their role in modelling disease severity. These data reflect fundamental inefficiencies in muscle formation, and sarcomeric force transmission, consistent with muscle atrophy. The most significant defects in both muscle structure and function were seen in the more severe FSHD individuals; this emphasises the need to include detailed genetic and clinical phenotyping of all individuals in pre-clinical research and drug discovery/therapeutic screening analyses to ensure a clear understanding of baseline disease pathology at the time of analysis.

Finally, we show that adenine base editing of the most severe FSHD individuals iPSC’s produced both a structural and functional improvements in disease pathology. In 2D, *DUX4* and its downstream targets (*ZSCAN4*, *TRIM43* and *PRAMEF1*) were all reduced toward healthy control levels during differentiation. While *DUX4* and *PRAMEF1* expression were no longer significantly different from healthy controls, both *ZSCAN4* and *TRIM43* remained higher in ABE 3D EMTs, at similar levels to that seen in mild FSHD. Morphologically, ABE improved 2D myofibre organization and increased myotube area, so that it was no longer different to healthy controls. In 3D, force generation was increased by over 600%, compared to the untreated FSHD 3D EMT and resulted in an improvement in the force–frequency profile, muscle mass and twitch-tetanus ratio; however phenotypic rescue in ABE 3D EMTs was not complete with function restored to that of a mild FSHD level. While complete correction of disease was not achieved, this data strengthens the rationale for precise genome editing strategies as a potential therapy for FSHD. The observation that functional gains accompany a reduction in the cytotoxic levels of *DUX4* (and downstream target genes) supports the use of DUX4-targeted gene therapies in FSHD to improve muscle function and performance. However, this approach alone did not result in the complete restoration of disease pathology and the need for combinatorial treatments that target both hypomethylation of the D4Z4 region and *DUX4* expression or promote muscle regeneration may be required.

Several aspects of this work will inform future studies. First, the pathological variability among individuals with FSHD highlights the unique genetic and epigenetic heterogeneity intrinsic to this disease. Expanding these studies to a larger number of childhood-onset FSHD individuals will provide further confidence in the use of iPSC-derived skeletal muscle models as a predictive tool to study disease pathobiology *in vitro*. Next, while ddPCR and immunofluorescence analyses provide robust and targeted readouts, a multiomics (transcript and proteome) analysis may capture the pattern of *DUX4* expression and potentially identify novel pathways responsible for disease severity in FSHD. Finally, this work highlights that targeted ABE of *DUX4* improves muscle function, however this therapeutic approach did not result in a function ‘cure’ which suggests that reducing *DUX4* mRNA expression alone, even at an ‘embryonic’ level as done here using iPSCs, is not sufficient, therefore combination therapy approaches may be required to restore muscle function and performance in FSHD. Furthermore, longitudinal analyses of gene editing approaches will aid in determining functional stability of this treatment, which will be critical given the chronic nature of FSHD.

## Conclusion

Across complementary 2 and 3D iPSC-derived skeletal muscle models, we identify severity-linked molecular, structural, and functional phenotypes that align with FSHD pathology. In 2D myotubes, *DUX4* and downstream target genes are induced early during differentiation, with the largest and most persistent activation in clinically severe individuals, accompanied by reductions in ACTN2+ myotube volume and area. In 3D EMTs, FSHD tissues retain the disease transcriptional signature and exhibit profound weakness in twitch and tetanic force generation, blunted force–frequency responses, reduced tissue mass, and markedly reduced specific force, again tracking with patient severity. Finally, targeted gene therapy using adenine base editing suppresses *DUX4* pathway activation and improves functional measures in the most severe FSHD individual, providing important proof-of-concept data in support of targeted genetic intervention in the mitigation of disease-associated phenotypes in FSHD.

## Methods

### Sex as a biological variable, study approval ethics and consent

Our study examined both male and female individuals, and similar findings are reported for both sexes. All participants provided informed consent through the Royal Children’s Hospitals (RCH) Human research ethics committee (HREC) as part of the pediatric <u>FSHD</u> <u>L</u>ongitudinal <u>O</u>utcomes <u>S</u>tudy (iFSHD-LOS, HREC 76964, ACTRN12621001293853).

### Statistical analyses

All statistical analyses were performed using Graphpad prism version 11.0.1 for Windows, GraphPad Software, Boston, Massachusetts USA, www.graphpad.com. Either one or two-way ANOVA’s with correction for multiple testing (Tukey) were performed as outlined in the figure legends. All data are shown as mean ± standard deviations (SD), unless otherwise stated in the figure legend.

### Clinical assessments, FSHD clinical score, FSHD COM Peds and 6min walk distance

The iFSHD-LOS was a natural history study which enrolled twenty-one children and young adults and followed disease progress over a two-year period between 2022 and 2024. All participants had a genetically confirmed diagnosis of FSHD type 1 and attended study visits at a single site, the RCH in Melbourne, Australia where they were assessed by the same team encompassing a doctor and a physiotherapist. At each visit, occurring at annual intervals, measures of muscle strength and function were assessed as well as blood taken for biobanking and iPSC generation. The clinical assessments performed included the FSHD clinical score (FCS), FSHD composite outcome measure for pediatrics (FSHD-COM Peds) and the six minute walk distance (6MWD).

The FCS measures facial, shoulder, arm, pelvic girdle, distal lower limb and trunk muscle strength with scores ranging from 0 to 15 [21]. A score of 1 indicating barely detectable weakness and 15 indicating severe weakness in all muscle groups. In this study a FCS score ranging from 1 to 5 was classified as mild, 6 to 10 moderate and 11 to 15 as severe.

FSHD-COM, an 18-item disease-specific assessment that evaluates leg, arm and trunk function, grip strength and functional balance, uses scores range from 1 to 84 points, where a higher score indicates greater functional limitation [22].

### Generation of induced pluripotent stem cell lines and validation of pluripotency

Four of the 21 participants (1 female (F): 3 male (M)) were selected for iPSCs derivation from peripheral blood monocytes (PBMCs). These individuals were selected as they were representative of the different stages of disease severity based on clinical scores and reflect the spectrum of disease phenotypes in FSHD.

iPSCs were generated by the MCRI’s iPSC Derivation and Gene Editing Facility using previously established methods [18].

Markers of pluripotency (OCT4, TRA-1–60, SSEA4, and TRA-1-81) were confirmed by antibody staining followed by flow cytometry. Briefly, iPSCs were dissociated with TrypLE (Thermo Fisher Scientific) and incubated with conjugated antibodies to cell surface proteins TRA-1–60, SSEA4, and TRA-1-81 (Table 2) diluted in PBS containing 2% fetal bovine serum (FBS) for 15 min at 4°C. iPSCs were washed with 2% FBS in PBS, fixed and permeabilized using the eBioscienceTM Foxp3/Transcription Factor Staining buffer set (Thermo Fischer Scientific), then stained with a conjugated antibody to intracellular OCT3/4 (Table 2). Single cells were analyzed using a LSR Fortessa X20 (BD Biosciences) and BD FACSDiva and FCS Express software.

**Table 2:**
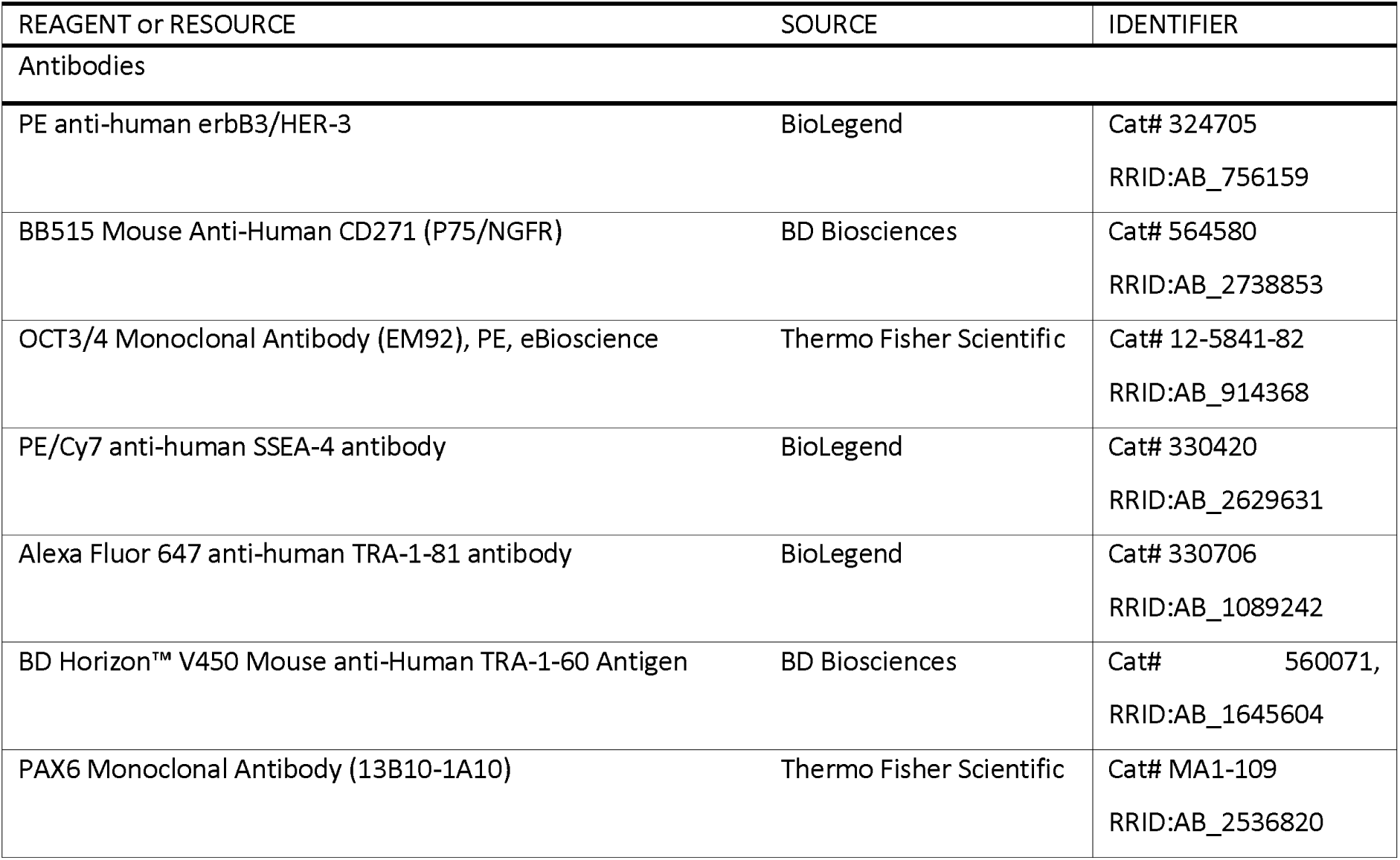
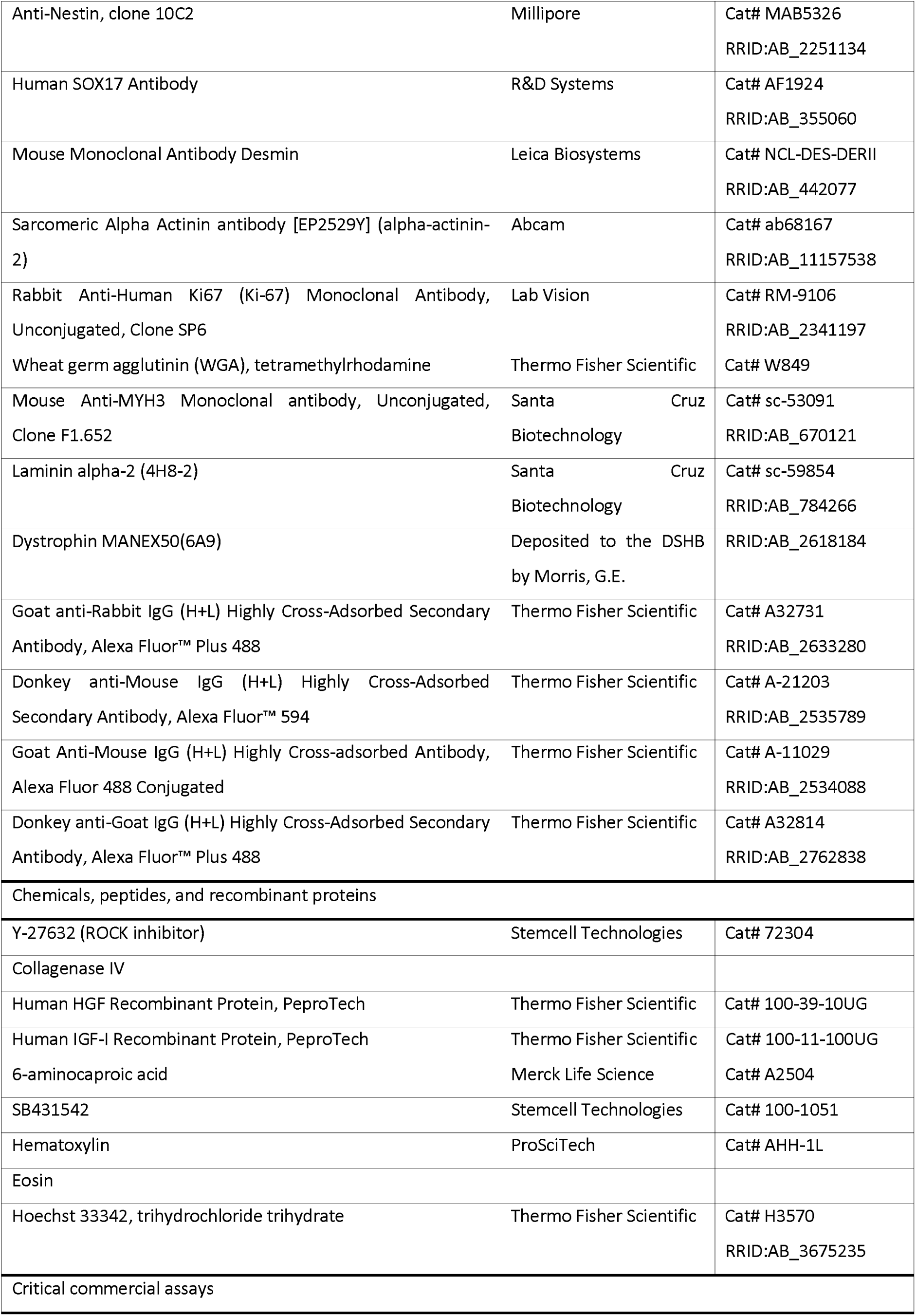
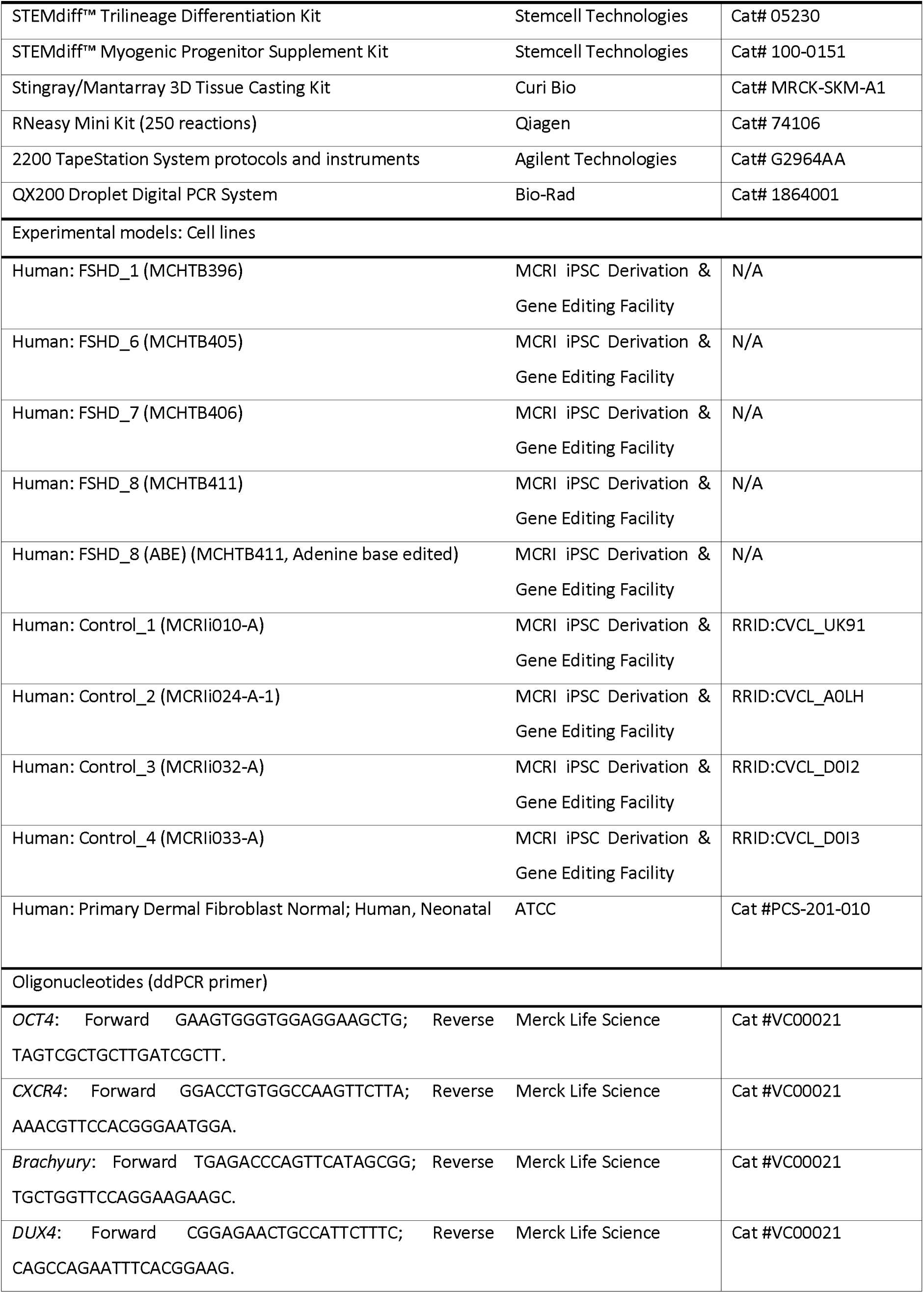
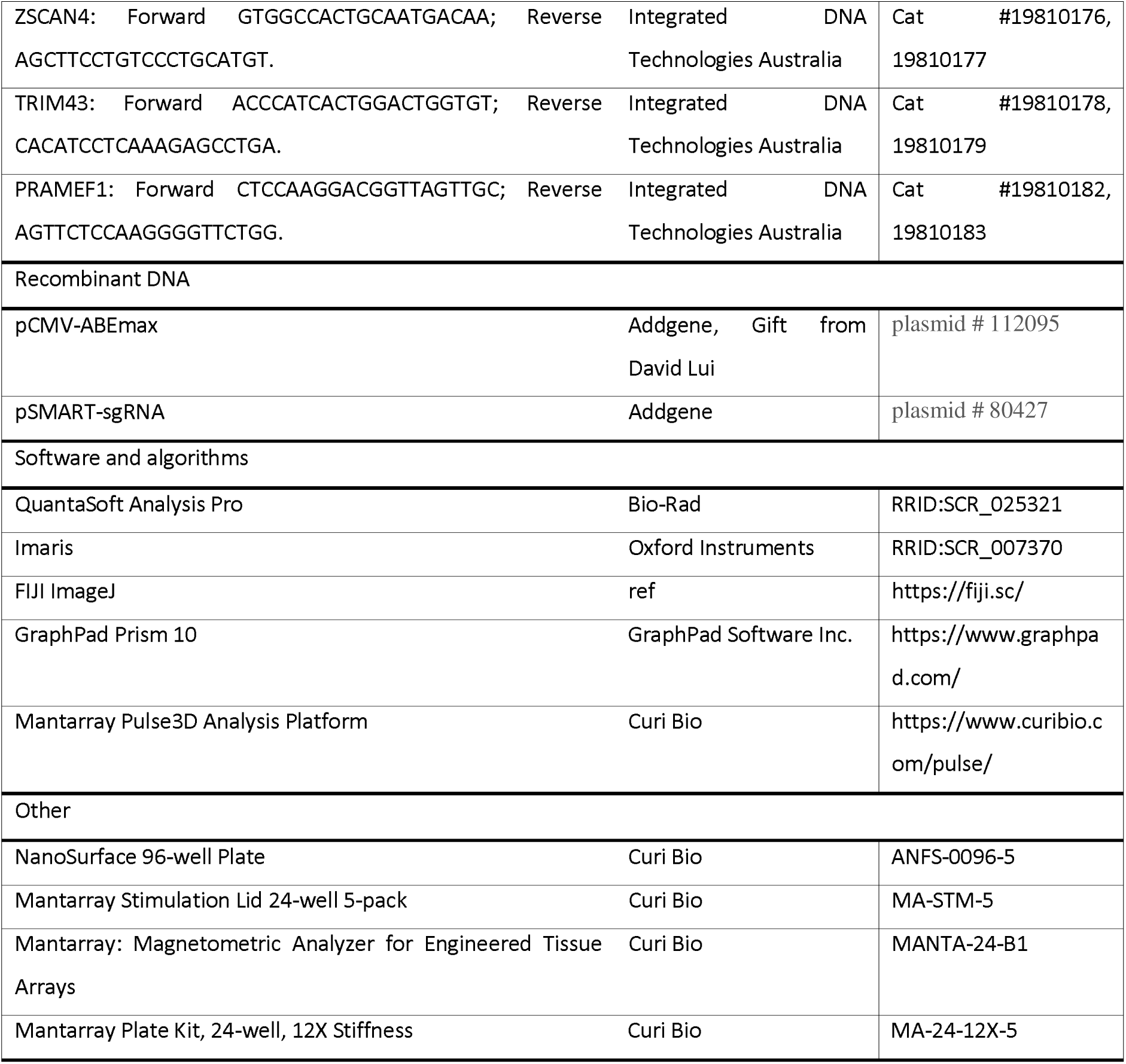
Key Resources.

iPSCs were validated for pluripotency by generating endoderm, mesoderm, and ectoderm using STEMdiff™ Trilineage Differentiation Kit trilineage differentiation kit (Stemcell Technologies) following manufacturer instructions. Endoderm and ectoderm markers were confirmed with immunofluorescent labelling (Table 2) while mesoderm markers were confirmed with ddPCR (Table 2).

### Base editing strategy for FSHD iPSCs

Plasmid encoding sgRNA specific to the *DUX4* PAS was generated by annealing oligonucleotides CACCGTTAAAATGCCCCCTCCCTG and AAACCAGGGAGGGGGCATTTTAAC (as outlined in [19]) followed by ligation into Bbs1-digested pSMART-sgRNA plasmid (Addgene plasmid # 80427). mRNA encoding ABEmax was generated from PmeI-digested pCMV-ABEmax (Addgene plasmid # 112095, a gift from David Lui) using the mMessage mMachine kit T7 Ultra kit (Thermo Fisher Scientific) according to the manufacturer’s recommendations. Base-editing factors (sgRNA plasmid and ABEmax mRNA) were introduced into iPSCs using the Neon transfection system (Thermo Fisher Scientific). Briefly, iPSCs were harvested using TryPLE and resuspended at a concentration of ∼10^7^ cells/ml in R buffer. ABEmax mRNA (2 µg), plasmid encoding DUX4-PAS-specific sgRNA (0.5 µg) and 10^5^ cells (10 µl) were assembled, aspirated into a 10 µl Neon tip and electroporated using the following conditions; 1100 V, 30 ms, 1 pulse. Cells were seeded onto Matrigel coated plate in E8 medium supplemented with CloneR2 (Stem Cell Technologies). Two rounds of base-editing was performed, with cells recovering for 72 hours before a subsequent round of electroporation.

### Cell culture maintenance

All cell lines were grown and maintained in a humidified incubator at 37°C and 5% CO_2_. iPSC lines and myogenic progenitors were grown on Matrigel (Corning)-coated plates.

iPSCs were cultured in mTeSR Plus complete medium (Stemcell Technologies), media was changed every 1-2 days, and cultures were passaged (1:4 – 1:10) every 3-4 days with 0.5 mM EDTA in DPBS (Gibco). For the first 24 hours post-thaw and post-passage, mTeSR Plus medium was supplemented with 10 μM ROCK inhibitor (Y-27632, Stemcell Technologies) to enhance cell survival.

Myogenic progenitors were cultured in Skeletal Muscle Cell Growth Medium-2 (SkGM-2, Lonza). Media was changed every 2-3 days, and myogenic progenitors were passaged (2500 cells/cm^2^) every 3-4 days with 1X TrypLE Select (Gibco).

### Cellular impedance assay

iPSCs were plated on Matrigel (Corning)-coated 96-well cellular impedance plates (Cat #300601010, xCELLigence, Agilent) at 10 000 cells per well. Cell impedances were monitored for 96 hours. Cell indices recorded using xCELLigence RTCA Software Pro (Agilent) were captured as an area under the curve (AUC) readout. Data were analyzed in GraphPad Prism using one-way ANOVA. Each datapoint represents the average of 6 wells.

### Generation of iPSC-derived myogenic progenitors

iPSC lines were differentiated into myogenic progenitors (myoblasts) using the STEMDiff™ Myogenic Progenitor Supplement Kit (Stemcell Technologies) and following manufacturer instructions. After 30 days, cells were dissociated using TrypLE Select (Gibco) and 1 mg/mL Collagenase IV (Sigma-Aldrich), filtered (70um), and resuspended as single cells in SkGM-2 (Lonza) supplemented with 10 μM ROCK inhibitor (Y-27632, Stemcell Technologies). Myogenic progenitors were plated onto Matrigel-coated tissue culture plates at 5,000 -15,000 cells/cm^2^. Media was changed to SkGM-2 without ROCK inhibitor after 24 hours.

### Terminal differentiation of myogenic progenitors to skeletal muscle myotubes in monolayer 2D format

All terminal differentiation experiments were performed using passage 4 or 5 myogenic progenitors. Myogenic progenitors were seeded in 24-well plates (Nunc) and grown to 90-100% confluency before being swapped into low-serum N2 media (DMEM/F12 [Gibco], 1% Glutamax [Gibco], 1% N2 supplement [Thermofisher], 1% Insulin-Transferrin-Selenium-Ethanolamine (ITS-X) [Thermofisher]). Media was changed every 2 days for 6 to 10 days as outlined in each figure. For volumetric analyses, myogenic progenitors were instead seeded into 96-well micropatterned etched plates (NanoSurface Plates, Curi Bio).

### Fluorescence-activated cell sorting (FACS)

Myogenic progenitors were dissociated with 1X TrypLE Select (Gibco) and resuspended in FACS buffer (1X PBS with 1 mg/mL bovine serum albumin and 2 mg/mL EDTA). Cells were stained with conjugated antibodies to the cell surface proteins ERBB3 and p75/NGFR (Table 2) for 15 min at 4°C. Dead cells were marked via Hoechst staining (Table 2). Myogenic progenitors were washed thrice in FACS buffer, passed through a 35 µm cell strainer, and sorted on a BD FACSAria Fusion cell sorter. Live single cells positive for ERBB3/NGFR were collected and expanded to passage 4 for 3D EMT generation.

### Generation of 3D engineered muscle tissues (EMT)

3D EMT were created according to manufacturer’s instructions (3D Tissue Casting Kit, Curi Bio [23]. ERBB3-positive myogenic progenitors and embryonic fibroblasts (human male neonate fibroblasts, passage 6, ATCPCS201010, ATCC) were suspended in a hydrogel mix containing Matrigel (Corning), thrombin (Thermo Fisher) and fibrinogen (Thermo Fisher). 3D EMTs were cultured in SkGM-2 containing 2 g/L 6-aminocaproic acid for the first 24 hours post-creation. On Day 0, 3D EMTs were transferred to fusion media [24], modified to a DMEM base [Thermo Fisher #10569010]). Media was changed every 2-3 days. From Day 10 of fusion onwards, the concentration of 6-aminocaproic acid was increased to 5 g/L.

### 3D EMT force measurements

3D EMTs were stimulated using a custom-built carbon electrode lid (Curi Bio). Electrical pulses were composed of biphasic signals of 5 ms per phase and 100 mA/-100 mA amplitude. Twitch kinetics were measured with three 1 Hz frequency pulses spaced 3 sec apart, and tetanus kinetics were measured with a single 50 Hz frequency pulses lasting 3 sec. Force frequency kinetics were captured through programmed 3 sec pulses with 30 sec rest between stimuli, and performed at 1 Hz, 2 Hz, 5 Hz, 10 Hz, 20 Hz, 35 Hz, 50 Hz, and 75 Hz. 3D EMTs were rested for 15 min between stimulation protocols to avoid fatigue. Concentric muscle contractions were measured by magnetometers built within the Mantarray platform [24]. Values were captured every 0.01 sec. Twitch and tetanus kinetics were measured daily from Day 7 of fusion onwards. Force frequency kinetics were conducted on either Day 14 or 24 as outlined in the figure legend.

Twitch, tetanus, and force frequency kinetics captured by the Mantarray system were analyzed in GraphPad Prism. The peak readout of each contraction was normalized by subtracting a baseline reading recorded 0.01 sec before each programmed stimulus began. Twitch:tetanus ratios were calculated by dividing twitch values by tetanus values recorded on the final day of culture (Day 14 or Day 24). Specific force was calculated by dividing the final 50Hz Tetanus reading by the weight of the matched muscle constructs (mg).

### RNA extraction and droplet digital quantitative real-time PCR (ddPCR) quantitation

RNA was extracted from cell cultures using the RNeasy mini kit (Qiagen). In brief, adherent iPSC or myogenic progenitor cells were washed in PBS, lysed with 350-600 µL RLT buffer, and homogenized using a 21-gauge needle and syringe. For 3D engineered muscle tissues, muscles were gently removed from lattice poles, blotted on lab bench roll, weighed, and snap frozen in liquid nitrogen. Tissues were lysed in 350 µL RLT buffer and homogenized using a T10 Basic Homogenizer (IKA). To remove cell debris, the homogenized tissues were centrifuged at maximum speed for 1 min, and the supernatant was transferred to a new 1.5 mL collection tube. For all samples, RNA was precipitated from solution using 70% ethanol and placed on a flow-through column with a silica membrane. RNA bound to the membrane was washed with provided buffers via microcentrifugation. RNA was further purified with on-column DNA digestion (DNase I, Qiagen) before being eluted in 30 μL MilliQ water. RNA quantity and quality was measured using TapeStation (Agilent Technologies 2200). Diluted RNA concentrations were assessed using Qubit 3.0 fluorometer (Thermo Fisher Scientific). Samples were diluted to 20 ng/μL, and 2 ng/µL of RNA were reverse transcribed to synthesise complementary DNA (cDNA) using the High-Capacity cDNA Reverse Transcription Kit (Thermo Fisher Scientific) as per manufacturer guidelines. In brief, RNA was combined with reverse transcription buffer, random primers, dNTPs, Multiscribe, RNase inhibitor, and MilliQ water for subsequent PCR reverse transcription at the following thermal cycling conditions: 10 min at 25°C, 60 min at 37°C, and 5 min at 95°C.

ddPCR assays were conducted using 2X QX200 ddPCR EvaGreen Supermix (Biorad) in a twin.tec 96-well plate (Biorad) to a final volume of 24 μl for lipid droplet generation. The sample plate of droplets was placed in a thermal cycler (T100, BioRad) for subsequent PCR amplification using gene-specific primers outlined in Table 2. The thermal cycling conditions were as follows: 1 activation cycle of 5 min at 95°C, 40 denaturation cycles of 30 sec at 96°C and annealing cycles of 1 min at 55–60°C depending on the target gene of interest, a post-cycling step of signal stabilisation of 5 min at 4°C followed by 5 min at 90°C. All cycling steps were performed using a 2°C per sec ramp rate. Following PCR amplification, the sample plate was loaded on the QX200 Droplet reader (Biorad) and the assay information was entered into QuantaSoft (BioRad) software.

### Cryosectioning

3D EMTs were removed from lattice poles, briefly blotted on tissue paper, weighed, and placed in Tissue-Tek Cryomolds (ProSciTech). The constructs were covered in Tissue-Tek OCT (Optimal Cutting Temperature) compound and submerged into liquid nitrogen-cooled isopentane. Cryopreserved 3D EMTs were stored at -80°C until 12 µm cross-sectional slices (Leica CM3050S Cryostat) were mounted on microscope slides (Superfrost Plus Slides, Epredia) and stored at -80°C until use.

### Immunofluorescence staining

iPSCs and myogenic progenitors were fixed in 4% paraformaldehyde in PBS for 20 min at room temperature, then permeabilized with 0.1% Triton X-100 in PBS for 10 min at room temperature. Non-specific binding was blocked with 3% bovine serum albumin in PBS containing 0.1% Tween-20 (PBS-T) overnight at 4°C. Cells were incubated for 2 hrs at room temperature with primary antibodies diluted in PBS-T containing 1% bovine serum albumin, followed by secondary antibodies for 1 hr (Table 2). Nuclei were stained with Hoechst (1 µg/mL, Thermo Fisher). Cells were washed thrice in PBS and imaged with a confocal microscope (Andor Dragonfly Spinning Disk Confocal 200, or Zeiss LSM 900).

3D EMT cryosections on microscope slides were left to thaw and air-dry at room temperature. All further steps were performed in a humidified dark chamber. All antibodies were diluted in 2% bovine serum albumin in 1X PBS, while 0.1% Triton X-100 in 1X PBS was used for triple washes between each incubation step. After 1 hr of non-specific blocking at room temperature, slides were incubated overnight at 4°C with 1:50 dilution of alpha-actinin-2 primary antibody (Table 2), incubated with donkey anti-rabbit IgG 488 1:500 for 1 hr at room temperature, in the dark, then stained with 1:100 wheat germ agglutinin– tetramethylrhodamine and 1:5000 Hoechst for 1 hr. Sections were fixed in 4% PFA for 1 hr before washed slides were mounted with Immumount (Epredia) and a coverslip. Slides were imaged on an Axio Imager.Z2 (Zeiss).

### Immunocytochemistry analyses (ImageJ, Imaris, Zen)

The approximate size of myotubes in monolayer 2D cultures was assessed by measuring alpha-actinin-2 -stained areas in immunofluorescent images. The areas of alpha-actinin-2 positive fluorescent staining were compared to the total area of each image and converted to a percentage ratio. One-way ANOVAs (GraphPad Prism) were used for statistical analysis.

Individual myotube length and volume analyses were conducted on myotubes grown on micropatterned plates (Curi Bio) as outlined above. A total of 65 z-stack fluorescent images taken at 0.25 µM steps per individual were loaded to Imaris. 3D objects were created from the fluorescent stained images. Imaris statistics automatically calculated internal volume and length, using the specific parameter BoundingBoxOO Length C. Images with overlapping myotubes were carefully 3D modelled within Imaris to separate cells. Volume and length data were analyzed using two-way ANOVAs in GraphPad Prism.

## Data availability

Supporting data including statistical analyses and reported means, confidence intervals and statistical outcomes are available in the supplemental material as outlined in the results and figure legends. All other data is available upon request.

## Author contributions

P.J.H and I.R.W conceived the project. P.J.H, V.G.C, L.K, K.M, C.T, C.C.C, C.H, N.T, R.J.M, S.E.H, K.dV contributed to the acquisition and/or analysis of data. S.E.H performed the adenine base editing. P.J.H, K.N.N and I.R.W contributed to the interpretation of data. P.J.H and I.R.W wrote the original manuscript. All authors reviewed and approved the submitted manuscript.

## Acknowledgements

We acknowledge the support provided by the Gillin Boys Foundation in acquiring the Curi Bio Mantarray platform. Services and technical assistance of the MCRI iPS cell and gene editing core facility were provided. This facility was established using a generous donation from the Stafford Fox Medical Research Foundation and is currently supported by Phenomics Australia and the Novo Nordisk Foundation reNEW Center for Stem Cell Medicine (NNF21CC0073729).

