## Supplementary figures and images for "Targeted *DUX4* base editing improves muscle function in an iPSC-derived model of childhood-onset FSHD"

### Supp Figure 1 C- H

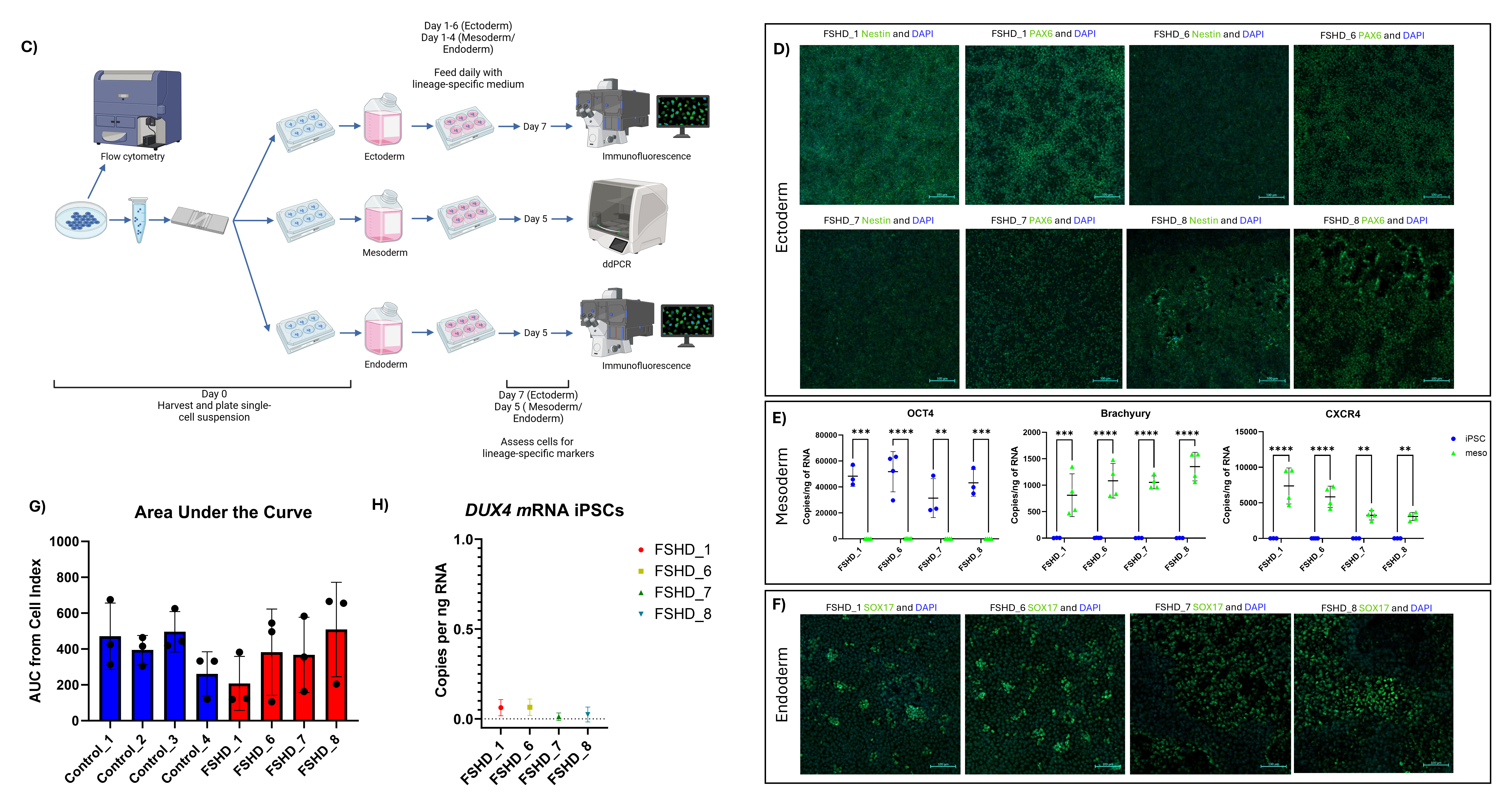

### Supp Figure 1A - B

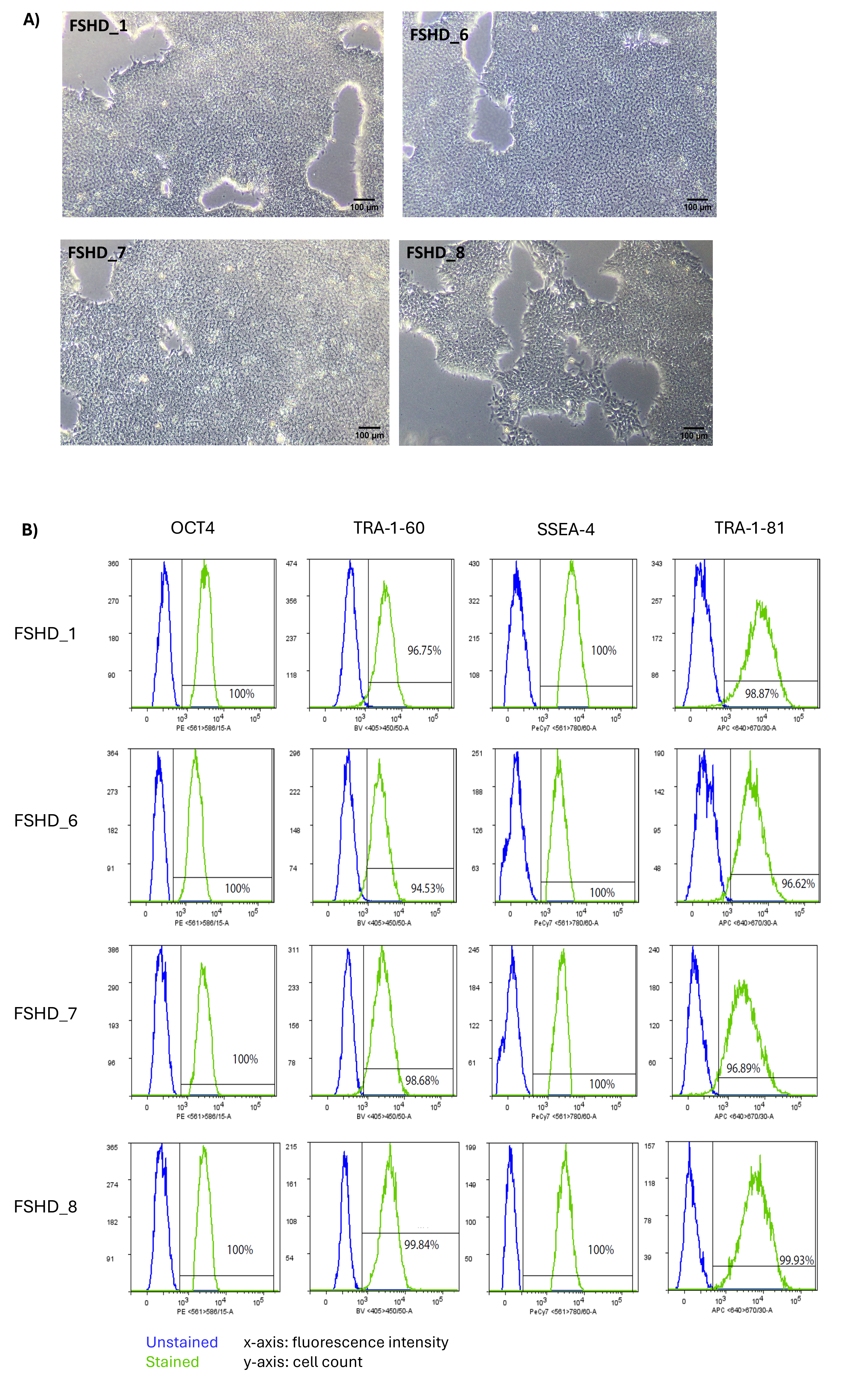
