## Supplementary material for "Targeted *DUX4* base editing improves muscle function in an iPSC-derived model of childhood-onset FSHD": Supp tables

**Supplementary Table 1**

**Two-way ANOVA statistical analysis summary of twitch force measures for figure 3F**

| Two-way ANOVA | Ordinary | |  |  |  | | |  |
| --- | --- | --- | --- | --- | --- | --- | --- | --- |
| Alpha | 0.05 | |  |  |  | | |  |
| Source of Variation | % of total variation | | P value | P value summary | Significant? | | |  |
| Interaction | 7.667 | | <0.0001 | **** | Yes | | |  |
| Time | 6.400 | | <0.0001 | **** | Yes | | |  |
| Individual | 73.06 | | <0.0001 | **** | Yes | | |  |
| ANOVA table | SS (Type III) | | DF | MS | F (DFn, DFd) | | | P value |
| Interaction | 1088731 | | 28 | 38883 | F (28, 344) = 10.08 | | | P<0.0001 |
| Time | 908783 | | 7 | 129826 | F (7, 344) = 33.65 | | | P<0.0001 |
| Individual | 10374935 | | 4 | 2593734 | F (4, 344) = 672.2 | | | P<0.0001 |
| Residual | 1327276 | | 344 | 3858 |  | | |  |
| Total | 14200573 | | 383 |  |  | | |  |
| **Tukey's multiple comparisons test** | | **Predicted (LS)**  **Mean diff.** | | **95.00% CI of diff.** | | **Summary** | **Adjusted P Value** | |
| D7 | |  | |  | |  |  | |
| Control_1 vs. Control_2 | | -82.62 | | -167.8 to 2.546 | | ns | 0.0621 | |
| Control_1 vs. FSHD_6 | | -171.9 | | -257.0 to -86.70 | | **** | <0.0001 | |
| Control_1 vs. FSHD_7 | | 328.9 | | 243.7 to 414.1 | | **** | <0.0001 | |
| Control_1 vs. FSHD_8 | | 409.4 | | 311.0 to 507.7 | | **** | <0.0001 | |
| Control_2 vs. FSHD_6 | | -89.25 | | -158.8 to -19.71 | | ** | 0.0044 | |
| Control_2 vs. FSHD_7 | | 411.5 | | 342.0 to 481.0 | | **** | <0.0001 | |
| Control_2 vs. FSHD_8 | | 492 | | 406.8 to 577.2 | | **** | <0.0001 | |
| FSHD_6 vs. FSHD_7 | | 500.8 | | 431.2 to 570.3 | | **** | <0.0001 | |
| FSHD_6 vs. FSHD_8 | | 581.2 | | 496.1 to 666.4 | | **** | <0.0001 | |
| FSHD_7 vs. FSHD_8 | | 80.48 | | -4.689 to 165.7 | | ns | 0.0742 | |
| D8 | |  | |  | |  |  | |
| Control_1 vs. Control_2 | | -168.3 | | -253.5 to -83.14 | | **** | <0.0001 | |
| Control_1 vs. FSHD_6 | | -59.45 | | -144.6 to 25.72 | | ns | 0.3118 | |
| Control_1 vs. FSHD_7 | | 287 | | 201.9 to 372.2 | | **** | <0.0001 | |
| Control_1 vs. FSHD_8 | | 329.9 | | 231.6 to 428.3 | | **** | <0.0001 | |
| Control_2 vs. FSHD_6 | | 108.9 | | 39.32 to 178.4 | | *** | 0.0002 | |
| Control_2 vs. FSHD_7 | | 455.3 | | 385.8 to 524.9 | | **** | <0.0001 | |
| Control_2 vs. FSHD_8 | | 498.2 | | 413.1 to 583.4 | | **** | <0.0001 | |
| FSHD_6 vs. FSHD_7 | | 346.5 | | 276.9 to 416.0 | | **** | <0.0001 | |
| FSHD_6 vs. FSHD_8 | | 389.4 | | 304.2 to 474.6 | | **** | <0.0001 | |
| FSHD_7 vs. FSHD_8 | | 42.9 | | -42.27 to 128.1 | | ns | 0.6401 | |
| D9 | |  | |  | |  |  | |
| Control_1 vs. Control_2 | | 134.4 | | 49.20 to 219.5 | | *** | 0.0002 | |
| Control_1 vs. FSHD_6 | | 261.2 | | 176.0 to 346.4 | | **** | <0.0001 | |
| Control_1 vs. FSHD_7 | | 509.3 | | 424.1 to 594.5 | | **** | <0.0001 | |
| Control_1 vs. FSHD_8 | | 546.6 | | 448.3 to 645.0 | | **** | <0.0001 | |
| Control_2 vs. FSHD_6 | | 126.8 | | 57.30 to 196.4 | | **** | <0.0001 | |
| Control_2 vs. FSHD_7 | | 374.9 | | 305.4 to 444.5 | | **** | <0.0001 | |
| Control_2 vs. FSHD_8 | | 412.3 | | 327.1 to 497.4 | | **** | <0.0001 | |
| FSHD_6 vs. FSHD_7 | | 248.1 | | 178.6 to 317.6 | | **** | <0.0001 | |
| FSHD_6 vs. FSHD_8 | | 285.4 | | 200.3 to 370.6 | | **** | <0.0001 | |
| FSHD_7 vs. FSHD_8 | | 37.34 | | -47.83 to 122.5 | | ns | 0.7501 | |
| D10 | |  | |  | |  |  | |
| Control_1 vs. Control_2 | | 45.75 | | -39.42 to 130.9 | | ns | 0.5807 | |
| Control_1 vs. FSHD_6 | | 98.24 | | 13.07 to 183.4 | | * | 0.0146 | |
| Control_1 vs. FSHD_7 | | 317.6 | | 232.5 to 402.8 | | **** | <0.0001 | |
| Control_1 vs. FSHD_8 | | 344 | | 245.7 to 442.4 | | **** | <0.0001 | |
| Control_2 vs. FSHD_6 | | 52.49 | | -17.05 to 122.0 | | ns | 0.2357 | |
| Control_2 vs. FSHD_7 | | 271.9 | | 202.3 to 341.4 | | **** | <0.0001 | |
| Control_2 vs. FSHD_8 | | 298.3 | | 213.1 to 383.4 | | **** | <0.0001 | |
| FSHD_6 vs. FSHD_7 | | 219.4 | | 149.9 to 288.9 | | **** | <0.0001 | |
| FSHD_6 vs. FSHD_8 | | 245.8 | | 160.6 to 330.9 | | **** | <0.0001 | |
| FSHD_7 vs. FSHD_8 | | 26.38 | | -58.79 to 111.5 | | ns | 0.9149 | |
| D11 | |  | |  | |  |  | |
| Control_1 vs. Control_2 | | 88.84 | | 3.666 to 174.0 | | * | 0.0361 | |
| Control_1 vs. FSHD_6 | | 144.8 | | 59.65 to 230.0 | | **** | <0.0001 | |
| Control_1 vs. FSHD_7 | | 393.1 | | 307.9 to 478.2 | | **** | <0.0001 | |
| Control_1 vs. FSHD_8 | | 442 | | 343.7 to 540.3 | | **** | <0.0001 | |
| Control_2 vs. FSHD_6 | | 55.98 | | -13.56 to 125.5 | | ns | 0.1793 | |
| Control_2 vs. FSHD_7 | | 304.2 | | 234.7 to 373.8 | | **** | <0.0001 | |
| Control_2 vs. FSHD_8 | | 353.2 | | 268.0 to 438.3 | | **** | <0.0001 | |
| FSHD_6 vs. FSHD_7 | | 248.2 | | 178.7 to 317.8 | | **** | <0.0001 | |
| FSHD_6 vs. FSHD_8 | | 297.2 | | 212.0 to 382.4 | | **** | <0.0001 | |
| FSHD_7 vs. FSHD_8 | | 48.94 | | -36.23 to 134.1 | | ns | 0.5139 | |
| D12 | |  | |  | |  |  | |
| Control_1 vs. Control_2 | | 63.72 | | -21.45 to 148.9 | | ns | 0.2438 | |
| Control_1 vs. FSHD_6 | | 153.1 | | 67.93 to 238.3 | | **** | <0.0001 | |
| Control_1 vs. FSHD_7 | | 334.2 | | 249.0 to 419.4 | | **** | <0.0001 | |
| Control_1 vs. FSHD_8 | | 377.4 | | 279.0 to 475.7 | | **** | <0.0001 | |
| Control_2 vs. FSHD_6 | | 89.38 | | 19.83 to 158.9 | | ** | 0.0044 | |
| Control_2 vs. FSHD_7 | | 270.5 | | 200.9 to 340.0 | | **** | <0.0001 | |
| Control_2 vs. FSHD_8 | | 313.7 | | 228.5 to 398.8 | | **** | <0.0001 | |
| FSHD_6 vs. FSHD_7 | | 181.1 | | 111.6 to 250.6 | | **** | <0.0001 | |
| FSHD_6 vs. FSHD_8 | | 224.3 | | 139.1 to 309.5 | | **** | <0.0001 | |
| FSHD_7 vs. FSHD_8 | | 43.19 | | -41.98 to 128.4 | | ns | 0.6342 | |
| D13 | |  | |  | |  |  | |
| Control_1 vs. Control_2 | | 97.83 | | 12.66 to 183.0 | | * | 0.0152 | |
| Control_1 vs. FSHD_6 | | 196.5 | | 111.4 to 281.7 | | **** | <0.0001 | |
| Control_1 vs. FSHD_7 | | 391.9 | | 306.7 to 477.0 | | **** | <0.0001 | |
| Control_1 vs. FSHD_8 | | 443.6 | | 345.3 to 542.0 | | **** | <0.0001 | |
| Control_2 vs. FSHD_6 | | 98.71 | | 29.17 to 168.3 | | ** | 0.0011 | |
| Control_2 vs. FSHD_7 | | 294 | | 224.5 to 363.6 | | **** | <0.0001 | |
| Control_2 vs. FSHD_8 | | 345.8 | | 260.6 to 431.0 | | **** | <0.0001 | |
| FSHD_6 vs. FSHD_7 | | 195.3 | | 125.8 to 264.9 | | **** | <0.0001 | |
| FSHD_6 vs. FSHD_8 | | 247.1 | | 161.9 to 332.2 | | **** | <0.0001 | |
| FSHD_7 vs. FSHD_8 | | 51.74 | | -33.43 to 136.9 | | ns | 0.4565 | |
| D14 | |  | |  | |  |  | |
| Control_1 vs. Control_2 | | 80.64 | | -4.533 to 165.8 | | ns | 0.0732 | |
| Control_1 vs. FSHD_6 | | 214.7 | | 129.5 to 299.9 | | **** | <0.0001 | |
| Control_1 vs. FSHD_7 | | 399 | | 313.8 to 484.1 | | **** | <0.0001 | |
| Control_1 vs. FSHD_8 | | 450.6 | | 352.2 to 548.9 | | **** | <0.0001 | |
| Control_2 vs. FSHD_6 | | 134.1 | | 64.53 to 203.6 | | **** | <0.0001 | |
| Control_2 vs. FSHD_7 | | 318.3 | | 248.8 to 387.9 | | **** | <0.0001 | |
| Control_2 vs. FSHD_8 | | 369.9 | | 284.7 to 455.1 | | **** | <0.0001 | |
| FSHD_6 vs. FSHD_7 | | 184.3 | | 114.7 to 253.8 | | **** | <0.0001 | |
| FSHD_6 vs. FSHD_8 | | 235.8 | | 150.7 to 321.0 | | **** | <0.0001 | |
| FSHD_7 vs. FSHD_8 | | 51.59 | | -33.58 to 136.8 | | ns | 0.4595 | |

**Supplementary Table 2**

**Two-way ANOVA statistical analysis summary of twitch force measures for figure 3G**

| Two-way ANOVA | Ordinary |  |  |  |  |
| --- | --- | --- | --- | --- | --- |
| Alpha | 0.05 |  |  |  |  |
| Source of Variation | % of total variation | P value | P value summary | Significant? |  |
| Interaction | 7.911 | <0.0001 | **** | Yes |  |
| Time | 6.304 | <0.0001 | **** | Yes |  |
| Individual | 73.40 | <0.0001 | **** | Yes |  |
| ANOVA table | SS (Type III) | DF | MS | F (DFn, DFd) | P value |
| Interaction | 3710690 | 28 | 132525 | F (28, 344) = 10.56 | P<0.0001 |
| Time | 2956743 | 7 | 422392 | F (7, 344) = 33.65 | P<0.0001 |
| Individual | 34430467 | 4 | 8607617 | F (4, 344) = 685.8 | P<0.0001 |
| Residual | 4317613 | 344 | 12551 |  |  |
| Total | 46905437 | 383 |  |  |  |

| **Tukey's multiple comparisons**  **test** | **Predicted (LS)**  **Mean diff.** | **95.00% CI**  **of diff.** | **Summary** | **Adjusted P Value** |
| --- | --- | --- | --- | --- |
| D7 |  |  |  |  |
| Control_1 vs. Control_2 | -32.22 | -185.8 to 121.4 | ns | 0.9786 |
| Control_1 vs. FSHD_6 | -154.7 | -308.4 to -1.129 | * | 0.0473 |
| Control_1 vs. FSHD_7 | 755.5 | 601.9 to 909.1 | **** | <0.0001 |
| Control_1 vs. FSHD_8 | 880.3 | 702.9 to 1058 | **** | <0.0001 |
| Control_2 vs. FSHD_6 | -122.5 | -247.9 to 2.899 | ns | 0.0592 |
| Control_2 vs. FSHD_7 | 787.7 | 662.3 to 913.1 | **** | <0.0001 |
| Control_2 vs. FSHD_8 | 912.5 | 758.9 to 1066 | **** | <0.0001 |
| FSHD_6 vs. FSHD_7 | 910.2 | 784.8 to 1036 | **** | <0.0001 |
| FSHD_6 vs. FSHD_8 | 1035 | 881.4 to 1189 | **** | <0.0001 |
| FSHD_7 vs. FSHD_8 | 124.8 | -28.81 to 278.4 | ns | 0.1719 |
| D8 |  |  |  |  |
| Control_1 vs. Control_2 | 47.68 | -105.9 to 201.3 | ns | 0.9142 |
| Control_1 vs. FSHD_6 | 177.4 | 23.78 to 331.0 | * | 0.0144 |
| Control_1 vs. FSHD_7 | 808.4 | 654.8 to 962.0 | **** | <0.0001 |
| Control_1 vs. FSHD_8 | 888.1 | 710.7 to 1066 | **** | <0.0001 |
| Control_2 vs. FSHD_6 | 129.7 | 4.282 to 255.1 | * | 0.0386 |
| Control_2 vs. FSHD_7 | 760.7 | 635.3 to 886.2 | **** | <0.0001 |
| Control_2 vs. FSHD_8 | 840.4 | 686.8 to 994.1 | **** | <0.0001 |
| FSHD_6 vs. FSHD_7 | 631 | 505.6 to 756.5 | **** | <0.0001 |
| FSHD_6 vs. FSHD_8 | 710.7 | 557.1 to 864.3 | **** | <0.0001 |
| FSHD_7 vs. FSHD_8 | 79.7 | -73.91 to 233.3 | ns | 0.6133 |
| D9 |  |  |  |  |
| Control_1 vs. Control_2 | 219.6 | 66.02 to 373.2 | ** | 0.001 |
| Control_1 vs. FSHD_6 | 464 | 310.4 to 617.6 | **** | <0.0001 |
| Control_1 vs. FSHD_7 | 794.6 | 641.0 to 948.2 | **** | <0.0001 |
| Control_1 vs. FSHD_8 | 869.5 | 692.2 to 1047 | **** | <0.0001 |
| Control_2 vs. FSHD_6 | 244.4 | 118.9 to 369.8 | **** | <0.0001 |
| Control_2 vs. FSHD_7 | 574.9 | 449.5 to 700.4 | **** | <0.0001 |
| Control_2 vs. FSHD_8 | 649.9 | 496.3 to 803.5 | **** | <0.0001 |
| FSHD_6 vs. FSHD_7 | 330.6 | 205.2 to 456.0 | **** | <0.0001 |
| FSHD_6 vs. FSHD_8 | 405.5 | 251.9 to 559.1 | **** | <0.0001 |
| FSHD_7 vs. FSHD_8 | 74.95 | -78.66 to 228.6 | ns | 0.6676 |
| D10 |  |  |  |  |
| Control_1 vs. Control_2 | 169.4 | 15.80 to 323.0 | * | 0.0224 |
| Control_1 vs. FSHD_6 | 333.5 | 179.9 to 487.1 | **** | <0.0001 |
| Control_1 vs. FSHD_7 | 609.3 | 455.6 to 762.9 | **** | <0.0001 |
| Control_1 vs. FSHD_8 | 671 | 493.6 to 848.3 | **** | <0.0001 |
| Control_2 vs. FSHD_6 | 164.1 | 38.70 to 289.5 | ** | 0.0035 |
| Control_2 vs. FSHD_7 | 439.8 | 314.4 to 565.3 | **** | <0.0001 |
| Control_2 vs. FSHD_8 | 501.5 | 347.9 to 655.2 | **** | <0.0001 |
| FSHD_6 vs. FSHD_7 | 275.7 | 150.3 to 401.1 | **** | <0.0001 |
| FSHD_6 vs. FSHD_8 | 337.4 | 183.8 to 491.0 | **** | <0.0001 |
| FSHD_7 vs. FSHD_8 | 61.7 | -91.91 to 215.3 | ns | 0.8057 |
| D11 |  |  |  |  |
| Control_1 vs. Control_2 | 229.1 | 75.49 to 382.7 | *** | 0.0005 |
| Control_1 vs. FSHD_6 | 414.8 | 261.1 to 568.4 | **** | <0.0001 |
| Control_1 vs. FSHD_7 | 715 | 561.4 to 868.6 | **** | <0.0001 |
| Control_1 vs. FSHD_8 | 816.2 | 638.8 to 993.6 | **** | <0.0001 |
| Control_2 vs. FSHD_6 | 185.6 | 60.22 to 311.1 | *** | 0.0006 |
| Control_2 vs. FSHD_7 | 485.9 | 360.5 to 611.4 | **** | <0.0001 |
| Control_2 vs. FSHD_8 | 587.1 | 433.5 to 740.7 | **** | <0.0001 |
| FSHD_6 vs. FSHD_7 | 300.3 | 174.9 to 425.7 | **** | <0.0001 |
| FSHD_6 vs. FSHD_8 | 401.5 | 247.9 to 555.1 | **** | <0.0001 |
| FSHD_7 vs. FSHD_8 | 101.2 | -52.42 to 254.8 | ns | 0.3715 |
| D12 |  |  |  |  |
| Control_1 vs. Control_2 | 189.7 | 36.08 to 343.3 | ** | 0.007 |
| Control_1 vs. FSHD_6 | 448.1 | 294.5 to 601.7 | **** | <0.0001 |
| Control_1 vs. FSHD_7 | 669.8 | 516.1 to 823.4 | **** | <0.0001 |
| Control_1 vs. FSHD_8 | 765.8 | 588.4 to 943.2 | **** | <0.0001 |
| Control_2 vs. FSHD_6 | 258.4 | 133.0 to 383.9 | **** | <0.0001 |
| Control_2 vs. FSHD_7 | 480.1 | 354.6 to 605.5 | **** | <0.0001 |
| Control_2 vs. FSHD_8 | 576.1 | 422.5 to 729.7 | **** | <0.0001 |
| FSHD_6 vs. FSHD_7 | 221.6 | 96.21 to 347.1 | **** | <0.0001 |
| FSHD_6 vs. FSHD_8 | 317.7 | 164.1 to 471.3 | **** | <0.0001 |
| FSHD_7 vs. FSHD_8 | 96.03 | -57.59 to 249.6 | ns | 0.4265 |
| D13 |  |  |  |  |
| Control_1 vs. Control_2 | 221.7 | 68.06 to 375.3 | *** | 0.0009 |
| Control_1 vs. FSHD_6 | 522.3 | 368.7 to 675.9 | **** | <0.0001 |
| Control_1 vs. FSHD_7 | 792.1 | 638.5 to 945.7 | **** | <0.0001 |
| Control_1 vs. FSHD_8 | 890.6 | 713.3 to 1068 | **** | <0.0001 |
| Control_2 vs. FSHD_6 | 300.6 | 175.2 to 426.0 | **** | <0.0001 |
| Control_2 vs. FSHD_7 | 570.5 | 445.0 to 695.9 | **** | <0.0001 |
| Control_2 vs. FSHD_8 | 669 | 515.3 to 822.6 | **** | <0.0001 |
| FSHD_6 vs. FSHD_7 | 269.8 | 144.4 to 395.3 | **** | <0.0001 |
| FSHD_6 vs. FSHD_8 | 368.3 | 214.7 to 522.0 | **** | <0.0001 |
| FSHD_7 vs. FSHD_8 | 98.5 | -55.12 to 252.1 | ns | 0.3998 |
| D14 |  |  |  |  |
| Control_1 vs. Control_2 | 172.5 | 18.86 to 326.1 | * | 0.0189 |
| Control_1 vs. FSHD_6 | 558.5 | 404.9 to 712.1 | **** | <0.0001 |
| Control_1 vs. FSHD_7 | 820.3 | 666.7 to 973.9 | **** | <0.0001 |
| Control_1 vs. FSHD_8 | 919.8 | 742.4 to 1097 | **** | <0.0001 |
| Control_2 vs. FSHD_6 | 386 | 260.6 to 511.4 | **** | <0.0001 |
| Control_2 vs. FSHD_7 | 647.8 | 522.4 to 773.2 | **** | <0.0001 |
| Control_2 vs. FSHD_8 | 747.3 | 593.7 to 900.9 | **** | <0.0001 |
| FSHD_6 vs. FSHD_7 | 261.8 | 136.4 to 387.2 | **** | <0.0001 |
| FSHD_6 vs. FSHD_8 | 361.3 | 207.7 to 514.9 | **** | <0.0001 |
| FSHD_7 vs. FSHD_8 | 99.49 | -54.13 to 253.1 | ns | 0.3893 |

**Supplementary Table 3**

**Two-way ANOVA statistical analysis summary of twitch force measures for figure 6F**

| Two-way ANOVA | Ordinary |  |  |  |  |
| --- | --- | --- | --- | --- | --- |
| Alpha | 0.05 |  |  |  |  |
| Source of Variation | % of total variation | P value | P value summary | Significant? |  |
| Interaction | 21.27 | <0.0001 | **** | Yes |  |
| Row Factor | 7.082 | <0.0001 | **** | Yes |  |
| Column Factor | 68.66 | <0.0001 | **** | Yes |  |
| ANOVA table | SS (Type III) | DF | MS | F (DFn, DFd) | P value |
| Interaction | 1209980 | 51 | 23725 | F (51, 356) = 29.30 | P<0.0001 |
| Row Factor | 402879 | 17 | 23699 | F (17, 356) = 29.27 | P<0.0001 |
| Column Factor | 3905959 | 3 | 1301986 | F (3, 356) = 1608 | P<0.0001 |
| Residual | 288232 | 356 | 809.6 |  |  |
| Total | 5688622 | 427 |  |  |  |

| Tukey's multiple comparisons test | Predicted (LS)  Mean diff. | 95.00% CI of diff. | Summary | Adjusted P Value |
| --- | --- | --- | --- | --- |
| D7 |  |  |  |  |
| Control_2 vs. FSHD (mild) | -72.81 | -115.2 to -30.41 | **** | <0.0001 |
| Control_2 vs. FSHD (severe) | 147.1 | 104.7 to 189.5 | **** | <0.0001 |
| Control_2 vs. FSHD (base edited) | 18.84 | -23.57 to 61.24 | ns | 0.6609 |
| FSHD (mild) vs. FSHD (severe) | 219.9 | 177.5 to 262.3 | **** | <0.0001 |
| FSHD (mild) vs. FSHD (base edited) | 91.65 | 49.25 to 134.1 | **** | <0.0001 |
| FSHD (severe) vs. FSHD (base edited) | -128.3 | -170.7 to -85.86 | **** | <0.0001 |
| D8 |  |  |  |  |
| Control_2 vs. FSHD (mild) | -138.9 | -181.3 to -96.46 | **** | <0.0001 |
| Control_2 vs. FSHD (severe) | -17.49 | -59.89 to 24.91 | ns | 0.7113 |
| Control_2 vs. FSHD (base edited) | -129.9 | -172.3 to -87.50 | **** | <0.0001 |
| FSHD (mild) vs. FSHD (severe) | 121.4 | 78.97 to 163.8 | **** | <0.0001 |
| FSHD (mild) vs. FSHD (base edited) | 8.967 | -33.44 to 51.37 | ns | 0.9476 |
| FSHD (severe) vs. FSHD (base edited) | -112.4 | -154.8 to -70.01 | **** | <0.0001 |
| D9 |  |  |  |  |
| Control_2 vs. FSHD (mild) | 37.3 | -5.102 to 79.71 | ns | 0.1069 |
| Control_2 vs. FSHD (severe) | 107.8 | 65.38 to 150.2 | **** | <0.0001 |
| Control_2 vs. FSHD (base edited) | -4.43 | -46.83 to 37.97 | ns | 0.9931 |
| FSHD (mild) vs. FSHD (severe) | 70.48 | 28.08 to 112.9 | *** | 0.0001 |
| FSHD (mild) vs. FSHD (base edited) | -41.73 | -84.14 to 0.6724 | ns | 0.0556 |
| FSHD (severe) vs. FSHD (base edited) | -112.2 | -154.6 to -69.81 | **** | <0.0001 |
| D10 |  |  |  |  |
| Control_2 vs. FSHD (mild) | 36.72 | -5.689 to 79.12 | ns | 0.1159 |
| Control_2 vs. FSHD (severe) | 118.2 | 75.81 to 160.6 | **** | <0.0001 |
| Control_2 vs. FSHD (base edited) | 1.506 | -40.90 to 43.91 | ns | 0.9997 |
| FSHD (mild) vs. FSHD (severe) | 81.5 | 39.10 to 123.9 | **** | <0.0001 |
| FSHD (mild) vs. FSHD (base edited) | -35.21 | -77.61 to 7.194 | ns | 0.1416 |
| FSHD (severe) vs. FSHD (base edited) | -116.7 | -159.1 to -74.31 | **** | <0.0001 |
| D11 |  |  |  |  |
| Control_2 vs. FSHD (mild) | 23.21 | -19.19 to 65.61 | ns | 0.4921 |
| Control_2 vs. FSHD (severe) | 183 | 140.6 to 225.4 | **** | <0.0001 |
| Control_2 vs. FSHD (base edited) | 46.08 | 3.673 to 88.48 | * | 0.0271 |
| FSHD (mild) vs. FSHD (severe) | 159.8 | 117.4 to 202.2 | **** | <0.0001 |
| FSHD (mild) vs. FSHD (base edited) | 22.87 | -19.54 to 65.27 | ns | 0.5052 |
| FSHD (severe) vs. FSHD (base edited) | -136.9 | -179.3 to -94.53 | **** | <0.0001 |
| D12 |  |  |  |  |
| Control_2 vs. FSHD (mild) | 43.89 | 1.487 to 86.30 | * | 0.0393 |
| Control_2 vs. FSHD (severe) | 197.8 | 155.4 to 240.2 | **** | <0.0001 |
| Control_2 vs. FSHD (base edited) | 80.14 | 37.74 to 122.5 | **** | <0.0001 |
| FSHD (mild) vs. FSHD (severe) | 153.9 | 111.5 to 196.4 | **** | <0.0001 |
| FSHD (mild) vs. FSHD (base edited) | 36.25 | -6.154 to 78.65 | ns | 0.1234 |
| FSHD (severe) vs. FSHD (base edited) | -117.7 | -160.1 to -75.29 | **** | <0.0001 |
| D13 |  |  |  |  |
| Control_2 vs. FSHD (mild) | 25.21 | -17.20 to 67.61 | ns | 0.4179 |
| Control_2 vs. FSHD (severe) | 200.2 | 157.8 to 242.6 | **** | <0.0001 |
| Control_2 vs. FSHD (base edited) | 66.74 | 24.34 to 109.1 | *** | 0.0003 |
| FSHD (mild) vs. FSHD (severe) | 175 | 132.6 to 217.4 | **** | <0.0001 |
| FSHD (mild) vs. FSHD (base edited) | 41.54 | -0.8691 to 83.94 | ns | 0.0573 |
| FSHD (severe) vs. FSHD (base edited) | -133.4 | -175.8 to -91.03 | **** | <0.0001 |
| D14 |  |  |  |  |
| Control_2 vs. FSHD (mild) | 38.64 | -3.766 to 81.04 | ns | 0.0885 |
| Control_2 vs. FSHD (severe) | 189 | 146.6 to 231.4 | **** | <0.0001 |
| Control_2 vs. FSHD (base edited) | 67.07 | 24.67 to 109.5 | *** | 0.0003 |
| FSHD (mild) vs. FSHD (severe) | 150.4 | 108.0 to 192.8 | **** | <0.0001 |
| FSHD (mild) vs. FSHD (base edited) | 28.43 | -13.97 to 70.84 | ns | 0.3093 |
| FSHD (severe) vs. FSHD (base edited) | -121.9 | -164.4 to -79.54 | **** | <0.0001 |
| D15 |  |  |  |  |
| Control_2 vs. FSHD (mild) | 36.77 | -5.638 to 79.17 | ns | 0.1151 |
| Control_2 vs. FSHD (severe) | 231.9 | 189.5 to 274.3 | **** | <0.0001 |
| Control_2 vs. FSHD (base edited) | 74.21 | 31.80 to 116.6 | **** | <0.0001 |
| FSHD (mild) vs. FSHD (severe) | 195.1 | 152.7 to 237.5 | **** | <0.0001 |
| FSHD (mild) vs. FSHD (base edited) | 37.44 | -4.964 to 79.84 | ns | 0.1049 |
| FSHD (severe) vs. FSHD (base edited) | -157.7 | -200.1 to -115.2 | **** | <0.0001 |
| D16 |  |  |  |  |
| Control_2 vs. FSHD (mild) | 54.2 | 11.79 to 96.60 | ** | 0.0059 |
| Control_2 vs. FSHD (severe) | 219.5 | 177.1 to 262.0 | **** | <0.0001 |
| Control_2 vs. FSHD (base edited) | 73.63 | 31.22 to 116.0 | **** | <0.0001 |
| FSHD (mild) vs. FSHD (severe) | 165.4 | 122.9 to 207.8 | **** | <0.0001 |
| FSHD (mild) vs. FSHD (base edited) | 19.43 | -22.97 to 61.84 | ns | 0.6381 |
| FSHD (severe) vs. FSHD (base edited) | -145.9 | -188.3 to -103.5 | **** | <0.0001 |
| D17 |  |  |  |  |
| Control_2 vs. FSHD (mild) | 112.5 | 70.13 to 154.9 | **** | <0.0001 |
| Control_2 vs. FSHD (severe) | 287.6 | 243.1 to 332.1 | **** | <0.0001 |
| Control_2 vs. FSHD (base edited) | 121.4 | 76.94 to 165.9 | **** | <0.0001 |
| FSHD (mild) vs. FSHD (severe) | 175 | 130.6 to 219.5 | **** | <0.0001 |
| FSHD (mild) vs. FSHD (base edited) | 8.877 | -35.60 to 53.35 | ns | 0.9554 |
| FSHD (severe) vs. FSHD (base edited) | -166.2 | -212.6 to -119.7 | **** | <0.0001 |
| D18 |  |  |  |  |
| Control_2 vs. FSHD (mild) | 129.8 | 87.42 to 172.2 | **** | <0.0001 |
| Control_2 vs. FSHD (severe) | 352.4 | 310.0 to 394.8 | **** | <0.0001 |
| Control_2 vs. FSHD (base edited) | 153.4 | 111.0 to 195.8 | **** | <0.0001 |
| FSHD (mild) vs. FSHD (severe) | 222.6 | 180.1 to 265.0 | **** | <0.0001 |
| FSHD (mild) vs. FSHD (base edited) | 23.58 | -18.83 to 65.98 | ns | 0.4783 |
| FSHD (severe) vs. FSHD (base edited) | -199 | -241.4 to -156.6 | **** | <0.0001 |
| D19 |  |  |  |  |
| Control_2 vs. FSHD (mild) | 164.9 | 122.5 to 207.3 | **** | <0.0001 |
| Control_2 vs. FSHD (severe) | 356 | 313.6 to 398.4 | **** | <0.0001 |
| Control_2 vs. FSHD (base edited) | 174 | 131.6 to 216.4 | **** | <0.0001 |
| FSHD (mild) vs. FSHD (severe) | 191.1 | 148.7 to 233.5 | **** | <0.0001 |
| FSHD (mild) vs. FSHD (base edited) | 9.115 | -33.29 to 51.52 | ns | 0.9452 |
| FSHD (severe) vs. FSHD (base edited) | -182 | -224.4 to -139.6 | **** | <0.0001 |
| D20 |  |  |  |  |
| Control_2 vs. FSHD (mild) | 159.8 | 117.4 to 202.2 | **** | <0.0001 |
| Control_2 vs. FSHD (severe) | 356.8 | 314.4 to 399.2 | **** | <0.0001 |
| Control_2 vs. FSHD (base edited) | 169.7 | 127.3 to 212.1 | **** | <0.0001 |
| FSHD (mild) vs. FSHD (severe) | 197 | 154.6 to 239.4 | **** | <0.0001 |
| FSHD (mild) vs. FSHD (base edited) | 9.887 | -32.52 to 52.29 | ns | 0.9314 |
| FSHD (severe) vs. FSHD (base edited) | -187.1 | -229.5 to -144.7 | **** | <0.0001 |
| D21 |  |  |  |  |
| Control_2 vs. FSHD (mild) | 183.2 | 140.8 to 225.6 | **** | <0.0001 |
| Control_2 vs. FSHD (severe) | 372.7 | 330.3 to 415.2 | **** | <0.0001 |
| Control_2 vs. FSHD (base edited) | 187.7 | 145.3 to 230.1 | **** | <0.0001 |
| FSHD (mild) vs. FSHD (severe) | 189.5 | 147.1 to 231.9 | **** | <0.0001 |
| FSHD (mild) vs. FSHD (base edited) | 4.467 | -37.94 to 46.87 | ns | 0.993 |
| FSHD (severe) vs. FSHD (base edited) | -185.1 | -227.5 to -142.6 | **** | <0.0001 |
| D22 |  |  |  |  |
| Control_2 vs. FSHD (mild) | 229.6 | 187.2 to 272.0 | **** | <0.0001 |
| Control_2 vs. FSHD (severe) | 456 | 413.6 to 498.4 | **** | <0.0001 |
| Control_2 vs. FSHD (base edited) | 230 | 187.6 to 272.5 | **** | <0.0001 |
| FSHD (mild) vs. FSHD (severe) | 226.3 | 183.9 to 268.7 | **** | <0.0001 |
| FSHD (mild) vs. FSHD (base edited) | 0.4199 | -41.98 to 42.82 | ns | >0.9999 |
| FSHD (severe) vs. FSHD (base edited) | -225.9 | -268.3 to -183.5 | **** | <0.0001 |
| D23 |  |  |  |  |
| Control_2 vs. FSHD (mild) | 263 | 218.5 to 307.4 | **** | <0.0001 |
| Control_2 vs. FSHD (severe) | 500.7 | 456.2 to 545.1 | **** | <0.0001 |
| Control_2 vs. FSHD (base edited) | 261.5 | 217.1 to 306.0 | **** | <0.0001 |
| FSHD (mild) vs. FSHD (severe) | 237.7 | 195.3 to 280.1 | **** | <0.0001 |
| FSHD (mild) vs. FSHD (base edited) | -1.433 | -43.84 to 40.97 | ns | 0.9998 |
| FSHD (severe) vs. FSHD (base edited) | -239.1 | -281.5 to -196.7 | **** | <0.0001 |
| D24 |  |  |  |  |
| Control_2 vs. FSHD (mild) | 275.9 | 231.4 to 320.4 | **** | <0.0001 |
| Control_2 vs. FSHD (severe) | 521.9 | 477.4 to 566.4 | **** | <0.0001 |
| Control_2 vs. FSHD (base edited) | 280 | 235.5 to 324.4 | **** | <0.0001 |
| FSHD (mild) vs. FSHD (severe) | 246 | 203.6 to 288.4 | **** | <0.0001 |
| FSHD (mild) vs. FSHD (base edited) | 4.062 | -38.34 to 46.47 | ns | 0.9947 |
| FSHD (severe) vs. FSHD (base edited) | -241.9 | -284.3 to -199.5 | **** | <0.0001 |

**Supplementary Table 4**

**Statistical analysis summary of tetanic force measures for Figure 6G**

| Two-way ANOVA | Ordinary |  |  |  |  |
| --- | --- | --- | --- | --- | --- |
| Alpha | 0.05 |  |  |  |  |
| Source of Variation | % of total variation | P value | P value summary | Significant? |  |
| Interaction | 24.99 | <0.0001 | **** | Yes |  |
| Time | 7.647 | <0.0001 | **** | Yes |  |
| Disease | 66.03 | <0.0001 | **** | Yes |  |
| ANOVA table | SS (Type III) | DF | MS | F (DFn, DFd) | P value |
| Interaction | 4812908 | 51 | 94371 | F (51, 356) = 43.19 | P<0.0001 |
| Time | 1472591 | 17 | 86623 | F (17, 356) = 39.65 | P<0.0001 |
| Disease | 12715846 | 3 | 4238615 | F (3, 356) = 1940 | P<0.0001 |
| Residual | 777813 | 356 | 2185 |  |  |

| Tukey's multiple comparisons test | Predicted  (LS) Mean diff. | 95.00% CI | Summary | Adjusted P Value |
| --- | --- | --- | --- | --- |
| D7 |  |  |  |  |
| Control_2 vs. FSHD (mild) | -239.1 | -308.8 to -169.5 | **** | <0.0001 |
| Control_2 vs. FSHD (severe) | 286.8 | 217.1 to 356.4 | **** | <0.0001 |
| Control_2 vs. FSHD (base edited) | 83.37 | 13.71 to 153.0 | * | 0.0116 |
| FSHD (mild) vs. FSHD (severe) | 525.9 | 456.2 to 595.5 | **** | <0.0001 |
| FSHD (mild) vs. FSHD (base edited) | 322.5 | 252.8 to 392.2 | **** | <0.0001 |
| FSHD (severe) vs. FSHD (base edited) | -203.4 | -273.0 to -133.7 | **** | <0.0001 |
| D8 |  |  |  |  |
| Control_2 vs. FSHD (mild) | -96 | -165.7 to -26.35 | ** | 0.0024 |
| Control_2 vs. FSHD (severe) | 157.1 | 87.46 to 226.8 | **** | <0.0001 |
| Control_2 vs. FSHD (base edited) | 1.564 | -68.09 to 71.22 | ns | >0.9999 |
| FSHD (mild) vs. FSHD (severe) | 253.1 | 183.5 to 322.8 | **** | <0.0001 |
| FSHD (mild) vs. FSHD (base edited) | 97.57 | 27.91 to 167.2 | ** | 0.0019 |
| FSHD (severe) vs. FSHD (base edited) | -155.6 | -225.2 to -85.89 | **** | <0.0001 |
| D9 |  |  |  |  |
| Control_2 vs. FSHD (mild) | 89.78 | 20.12 to 159.4 | ** | 0.0053 |
| Control_2 vs. FSHD (severe) | 202.3 | 132.6 to 271.9 | **** | <0.0001 |
| Control_2 vs. FSHD (base edited) | 65.91 | -3.746 to 135.6 | ns | 0.0711 |
| FSHD (mild) vs. FSHD (severe) | 112.5 | 42.84 to 182.2 | *** | 0.0002 |
| FSHD (mild) vs. FSHD (base edited) | -23.87 | -93.53 to 45.79 | ns | 0.8129 |
| FSHD (severe) vs. FSHD (base edited) | -136.4 | -206.0 to -66.71 | **** | <0.0001 |
| D10 |  |  |  |  |
| Control_2 vs. FSHD (mild) | 94.86 | 25.20 to 164.5 | ** | 0.0028 |
| Control_2 vs. FSHD (severe) | 191.2 | 121.5 to 260.8 | **** | <0.0001 |
| Control_2 vs. FSHD (base edited) | 53.76 | -15.90 to 123.4 | ns | 0.1928 |
| FSHD (mild) vs. FSHD (severe) | 96.29 | 26.64 to 166.0 | ** | 0.0023 |
| FSHD (mild) vs. FSHD (base edited) | -41.09 | -110.8 to 28.56 | ns | 0.4249 |
| FSHD (severe) vs. FSHD (base edited) | -137.4 | -207.0 to -67.73 | **** | <0.0001 |
| D11 |  |  |  |  |
| Control_2 vs. FSHD (mild) | 125.5 | 55.87 to 195.2 | **** | <0.0001 |
| Control_2 vs. FSHD (severe) | 275.2 | 205.5 to 344.8 | **** | <0.0001 |
| Control_2 vs. FSHD (base edited) | 94.78 | 25.12 to 164.4 | ** | 0.0028 |
| FSHD (mild) vs. FSHD (severe) | 149.6 | 79.97 to 219.3 | **** | <0.0001 |
| FSHD (mild) vs. FSHD (base edited) | -30.75 | -100.4 to 38.91 | ns | 0.6653 |
| FSHD (severe) vs. FSHD (base edited) | -180.4 | -250.0 to -110.7 | **** | <0.0001 |
| D12 |  |  |  |  |
| Control_2 vs. FSHD (mild) | 132.4 | 62.70 to 202.0 | **** | <0.0001 |
| Control_2 vs. FSHD (severe) | 284.4 | 214.7 to 354.1 | **** | <0.0001 |
| Control_2 vs. FSHD (base edited) | 116.2 | 46.55 to 185.9 | *** | 0.0001 |
| FSHD (mild) vs. FSHD (severe) | 152 | 82.37 to 221.7 | **** | <0.0001 |
| FSHD (mild) vs. FSHD (base edited) | -16.16 | -85.82 to 53.50 | ns | 0.9324 |
| FSHD (severe) vs. FSHD (base edited) | -168.2 | -237.8 to -98.53 | **** | <0.0001 |
| D13 |  |  |  |  |
| Control_2 vs. FSHD (mild) | 121.7 | 52.07 to 191.4 | **** | <0.0001 |
| Control_2 vs. FSHD (severe) | 294.3 | 224.6 to 363.9 | **** | <0.0001 |
| Control_2 vs. FSHD (base edited) | 110.2 | 40.56 to 179.9 | *** | 0.0003 |
| FSHD (mild) vs. FSHD (severe) | 172.5 | 102.9 to 242.2 | **** | <0.0001 |
| FSHD (mild) vs. FSHD (base edited) | -11.51 | -81.17 to 58.15 | ns | 0.9739 |
| FSHD (severe) vs. FSHD (base edited) | -184 | -253.7 to -114.4 | **** | <0.0001 |
| D14 |  |  |  |  |
| Control_2 vs. FSHD (mild) | 154.4 | 84.70 to 224.0 | **** | <0.0001 |
| Control_2 vs. FSHD (severe) | 296.6 | 226.9 to 366.2 | **** | <0.0001 |
| Control_2 vs. FSHD (base edited) | 138.5 | 68.83 to 208.1 | **** | <0.0001 |
| FSHD (mild) vs. FSHD (severe) | 142.2 | 72.57 to 211.9 | **** | <0.0001 |
| FSHD (mild) vs. FSHD (base edited) | -15.87 | -85.53 to 53.79 | ns | 0.9356 |
| FSHD (severe) vs. FSHD (base edited) | -158.1 | -227.8 to -88.44 | **** | <0.0001 |
| D15 |  |  |  |  |
| Control_2 vs. FSHD (mild) | 160.2 | 90.55 to 229.9 | **** | <0.0001 |
| Control_2 vs. FSHD (severe) | 368.6 | 298.9 to 438.3 | **** | <0.0001 |
| Control_2 vs. FSHD (base edited) | 142.6 | 72.91 to 212.2 | **** | <0.0001 |
| FSHD (mild) vs. FSHD (severe) | 208.4 | 138.7 to 278.1 | **** | <0.0001 |
| FSHD (mild) vs. FSHD (base edited) | -17.64 | -87.30 to 52.02 | ns | 0.9142 |
| FSHD (severe) vs. FSHD (base edited) | -226 | -295.7 to -156.4 | **** | <0.0001 |
| D16 |  |  |  |  |
| Control_2 vs. FSHD (mild) | 222.5 | 152.9 to 292.2 | **** | <0.0001 |
| Control_2 vs. FSHD (severe) | 401.7 | 332.1 to 471.4 | **** | <0.0001 |
| Control_2 vs. FSHD (base edited) | 192.7 | 123.0 to 262.4 | **** | <0.0001 |
| FSHD (mild) vs. FSHD (severe) | 179.2 | 109.5 to 248.9 | **** | <0.0001 |
| FSHD (mild) vs. FSHD (base edited) | -29.81 | -99.47 to 39.85 | ns | 0.6869 |
| FSHD (severe) vs. FSHD (base edited) | -209 | -278.7 to -139.4 | **** | <0.0001 |
| D17 |  |  |  |  |
| Control_2 vs. FSHD (mild) | 300.4 | 230.8 to 370.1 | **** | <0.0001 |
| Control_2 vs. FSHD (severe) | 497.7 | 424.7 to 570.8 | **** | <0.0001 |
| Control_2 vs. FSHD (base edited) | 241.7 | 168.6 to 314.7 | **** | <0.0001 |
| FSHD (mild) vs. FSHD (severe) | 197.3 | 124.2 to 270.4 | **** | <0.0001 |
| FSHD (mild) vs. FSHD (base edited) | -58.78 | -131.8 to 14.28 | ns | 0.1627 |
| FSHD (severe) vs. FSHD (base edited) | -256.1 | -332.4 to -179.8 | **** | <0.0001 |
| D18 |  |  |  |  |
| Control_2 vs. FSHD (mild) | 356.1 | 286.5 to 425.8 | **** | <0.0001 |
| Control_2 vs. FSHD (severe) | 629.2 | 559.5 to 698.8 | **** | <0.0001 |
| Control_2 vs. FSHD (base edited) | 324 | 254.3 to 393.6 | **** | <0.0001 |
| FSHD (mild) vs. FSHD (severe) | 273 | 203.4 to 342.7 | **** | <0.0001 |
| FSHD (mild) vs. FSHD (base edited) | -32.19 | -101.8 to 37.47 | ns | 0.6318 |
| FSHD (severe) vs. FSHD (base edited) | -305.2 | -374.9 to -235.6 | **** | <0.0001 |
| D19 |  |  |  |  |
| Control_2 vs. FSHD (mild) | 397.8 | 328.2 to 467.5 | **** | <0.0001 |
| Control_2 vs. FSHD (severe) | 640 | 570.3 to 709.7 | **** | <0.0001 |
| Control_2 vs. FSHD (base edited) | 358.1 | 288.5 to 427.8 | **** | <0.0001 |
| FSHD (mild) vs. FSHD (severe) | 242.2 | 172.5 to 311.8 | **** | <0.0001 |
| FSHD (mild) vs. FSHD (base edited) | -39.68 | -109.3 to 29.98 | ns | 0.4567 |
| FSHD (severe) vs. FSHD (base edited) | -281.9 | -351.5 to -212.2 | **** | <0.0001 |
| D20 |  |  |  |  |
| Control_2 vs. FSHD (mild) | 396.6 | 326.9 to 466.3 | **** | <0.0001 |
| Control_2 vs. FSHD (severe) | 643.8 | 574.2 to 713.5 | **** | <0.0001 |
| Control_2 vs. FSHD (base edited) | 361.4 | 291.8 to 431.1 | **** | <0.0001 |
| FSHD (mild) vs. FSHD (severe) | 247.3 | 177.6 to 316.9 | **** | <0.0001 |
| FSHD (mild) vs. FSHD (base edited) | -35.17 | -104.8 to 34.49 | ns | 0.5615 |
| FSHD (severe) vs. FSHD (base edited) | -282.4 | -352.1 to -212.8 | **** | <0.0001 |
| D21 |  |  |  |  |
| Control_2 vs. FSHD (mild) | 488.3 | 418.6 to 558.0 | **** | <0.0001 |
| Control_2 vs. FSHD (severe) | 757.5 | 687.9 to 827.2 | **** | <0.0001 |
| Control_2 vs. FSHD (base edited) | 434.7 | 365.0 to 504.3 | **** | <0.0001 |
| FSHD (mild) vs. FSHD (severe) | 269.2 | 199.6 to 338.9 | **** | <0.0001 |
| FSHD (mild) vs. FSHD (base edited) | -53.62 | -123.3 to 16.04 | ns | 0.1948 |
| FSHD (severe) vs. FSHD (base edited) | -322.9 | -392.5 to -253.2 | **** | <0.0001 |
| D22 |  |  |  |  |
| Control_2 vs. FSHD (mild) | 558.1 | 488.5 to 627.8 | **** | <0.0001 |
| Control_2 vs. FSHD (severe) | 905.2 | 835.5 to 974.9 | **** | <0.0001 |
| Control_2 vs. FSHD (base edited) | 500.1 | 430.4 to 569.7 | **** | <0.0001 |
| FSHD (mild) vs. FSHD (severe) | 347.1 | 277.4 to 416.7 | **** | <0.0001 |
| FSHD (mild) vs. FSHD (base edited) | -58.03 | -127.7 to 11.62 | ns | 0.1394 |
| FSHD (severe) vs. FSHD (base edited) | -405.1 | -474.8 to -335.5 | **** | <0.0001 |
| D23 |  |  |  |  |
| Control_2 vs. FSHD (mild) | 594.4 | 521.3 to 667.4 | **** | <0.0001 |
| Control_2 vs. FSHD (severe) | 964.2 | 891.1 to 1037 | **** | <0.0001 |
| Control_2 vs. FSHD (base edited) | 554.9 | 481.9 to 628.0 | **** | <0.0001 |
| FSHD (mild) vs. FSHD (severe) | 369.8 | 300.2 to 439.5 | **** | <0.0001 |
| FSHD (mild) vs. FSHD (base edited) | -39.43 | -109.1 to 30.23 | ns | 0.4623 |
| FSHD (severe) vs. FSHD (base edited) | -409.3 | -478.9 to -339.6 | **** | <0.0001 |
| D24 |  |  |  |  |
| Control_2 vs. FSHD (mild) | 621.5 | 548.4 to 694.5 | **** | <0.0001 |
| Control_2 vs. FSHD (severe) | 1011 | 937.7 to 1084 | **** | <0.0001 |
| Control_2 vs. FSHD (base edited) | 592.5 | 519.4 to 665.5 | **** | <0.0001 |
| FSHD (mild) vs. FSHD (severe) | 389.2 | 319.6 to 458.9 | **** | <0.0001 |
| FSHD (mild) vs. FSHD (base edited) | -29.02 | -98.68 to 40.63 | ns | 0.7047 |
| FSHD (severe) vs. FSHD (base edited) | -418.3 | -487.9 to -348.6 | **** | <0.0001 |
